# Revisiting the *in vivo* functions of coilin in mice: Requirement for snRNA modifications and formation of novel condensates in elongating spermatids

**DOI:** 10.1101/2025.11.30.691380

**Authors:** Miyu Ichii, Yuma Ishigami, Xiao Yao, Jianxi Li, Josei Sato, Yuki Okada, Atsushi P. Kimura, Mari Mito, Shintaro Iwasaki, Yasuyuki Kurihara, Taichi Noda, Kimi Araki, Masato Ohtsuka, Saki Ohazama, Hiroshi Maita, Tsutomu Suzuki, Shinichi Nakagawa

## Abstract

Cajal bodies (CBs), first identified in the early 20th century, are classic nuclear structures organized primarily by the protein coilin. Although CBs have long been implicated in snRNA modification and maturation, their precise roles in mice remain unclear. Here, we revisit the functions of CBs by using genome editing to generate mice that lack nearly the entire coilin protein, retaining only the first seven amino acids. Mass-spectrometry analyses showed reduced 2′-*O*-methylation of U2 snRNA in coilin mutant liver; however, these reductions had minimal impact on overall gene expression and pre-mRNA splicing patterns. Additionally, detailed analysis recapitulated previously reported testicular size reductions and newly revealed decreased proportions of high-quality sperm. We also identified novel coilin-containing condensates distinct from genuine CBs in the nuclear pocket of elongating spermatids. Our findings implicate that coilin orchestrates membraneless organelles in various compartments, playing multifunctional roles beyond classical CB formation.

## Introduction

The nucleus is highly compartmentalized and contains various nuclear bodies that house proteins and nucleic acids involved in distinct nuclear processes, including nucleoli, nuclear speckles, paraspeckles, and others (Fox et al. 2018; Lafontaine et al. 2021; Ilik and Aktas 2022; Hirose et al. 2023). Also referred to as molecular condensates or membraneless organelles, these structures promote efficient molecular reactions by increasing the local concentration of specific factors and by segregating processes that could otherwise interfere with each other (Shin and Brangwynne 2017; Alberti and Hyman 2021). Nuclear bodies typically consist of defined sets of RNAs and RNA-binding proteins, many of which contain intrinsically disordered regions. Multivalent weak interactions among these components can drive liquid-liquid phase separation, resulting in condensate formation (Strom and Brangwynne 2019; Sabari et al. 2020; Roden and Gladfelter 2021).

Cajal bodies (CBs), first described by Ramón y Cajal and subsequently termed coiled bodies due to their distinct electron microscopic appearance, were molecularly identified through an autoantigen that specifically localized to these structures (Gall 2000). This autoantigen, termed coilin (Raska et al. 1991), was later confirmed to be an essential structural component necessary for CB formation. CBs have long been proposed to function primarily in the processing and assembly of small nuclear RNAs (snRNAs) into small nuclear ribonucleoprotein particles (snRNPs), critical constituents of the spliceosome required for pre-mRNA splicing (Machyna et al. 2015; Neugebauer 2017). Newly transcribed snRNAs are exported to the cytoplasm for Sm core protein assembly, then re-imported into the nucleus, where their final maturation, including 2′-*O*-methylation and pseudouridylation, occurs (Matera and Wang 2014). CBs are enriched in essential factors involved in snRNA modifications, such as the 2′-*O*-methyltransferase fibrillarin and scaRNAs, which guide these enzymes to specific target nucleotides within snRNAs. These observations support a functional role for CBs in the efficient maturation of snRNAs and snRNP biogenesis (Machyna et al. 2015; Meier 2017; Stanek 2017). Coilin plays a central architectural role in CB assembly, through multivalent interactions with key components such as SMN and NOPP140, interactions that are often mediated by methylated arginine residues (Courchaine et al. 2021; Courchaine et al. 2022; Ohazama et al. 2024; Arias Escayola et al. 2025).

Despite the identification of CBs over 120 years ago, their physiological significance remains somewhat enigmatic (Neugebauer 2017; Stanek 2023). Previous studies have shown that dependence on coilin for CB assembly and function varies considerably across different species. In *Drosophila melanogaster*, coilin-null flies are fully viable and fertile, displaying no obvious developmental abnormalities (Liu et al. 2009). Similarly, coilin mutant *Arabidopsis thaliana* plants exhibit normal growth under standard laboratory conditions but display compromised defense responses upon pathogen challenge, accompanied by increased intron retention in immunity-related genes (Collier et al. 2006; Abulfaraj et al. 2022). Coilin knockout mice exhibit partial embryonic lethality occurring between embryonic day (E)13.5 and birth; however, those mice that survive to adulthood appear morphologically normal without overt defects (Tucker et al. 2001; Walker et al. 2009). Consistent with these observations, disruption of coilin by genetic knockout or targeted mutations disperses CB components into the nucleoplasm; nonetheless, snRNA modifications continue to occur outside CBs in coilin mutants of *Drosophila* and mouse embryonic fibroblasts (Jady et al. 2003; Deryusheva and Gall 2009). The most severe phenotype associated with coilin depletion has been reported in zebrafish embryos, where coilin deficiency results in severe developmental arrest and lethality. Importantly, this phenotype can be rescued by supplementing mature snRNPs, but not fully modified snRNAs alone, indicating coilin’s critical role in accelerating snRNP assembly, particularly in rapidly developing organisms (Strzelecka et al. 2010).

Recent advancements in genome editing technologies, such as the iGONAD method (Gurumurthy et al. 2019), have enabled simpler and more efficient generation of mutant animals without time-consuming vector construction or reproductive manipulation. We thus revisited the physiological function of coilin by generating genome-edited mice with a more extensive deletion of the coilin gene than previous models, which retained the first 90 amino acids essential for coilin dimerization (Tucker et al. 2001). Consistent with earlier reports, our newly generated coilin mutant mice (*Coil^7aa^*/*Coil^7aa^*) were viable and exhibited no overt morphological defects. However, detailed analyses using mass spectrometry revealed that 2′-*O*-methylation of U2 snRNA was reduced in the liver tissue of these mutants. Furthermore, we observed a decrease in the proportion of high-quality sperm, despite grossly normal male fertility. Intriguingly, detailed analyses of testis tissue demonstrated that coilin transiently formed distinct molecular condensates in the "nuclear pocket" of elongating spermatids, structures clearly distinguishable from canonical nuclear CBs. These observations highlight coilin’s multifunctional potential in organizing membraneless organelles across multiple cellular compartments.

## Results

### Generation and validation of complete coilin knockout mice

Previous studies on coilin knockout mice (hereafter termed *Coil^82aa^*) retained the N-terminal 82 amino acids of coilin, a region required for coilin self-association. To further investigate the physiological functions of coilin and CBs, we generated genome-edited mice lacking nearly the entire coilin open reading frame, retaining only the first seven amino acids (MSKMAAS) and three amino acids (VAD) derived from the linker sequences (**Figure 1A-C**). Consistent with prior studies, we successfully obtained viable homozygous *Coil^7aa^*/*Coil^7aa^*adult mice by crossing heterozygous mice (see below). To verify the absence of CBs in *Coil^7aa^* mutants, we examined the localization of coilin and SMN, a protein known to co-localize with coilin in CBs, particularly in cortical neurons where prominent CBs are observed. In wildtype (WT) mice, coilin and SMN co-localized within distinct nuclear foci in approximately 10% of cortical neurons. In contrast, *Coil^7aa^*mutant neurons lacked nuclear SMN-positive foci (**Figure 1D**), indicating the complete absence of canonical CBs. We also examined the formation of previously reported residual CBs in *Coil^82aa^* mice, characterized by small, round nuclear foci containing Nopp140 but lacking the nucleolar-specific protein nucleophosmin 1 (Npm1). Due to limitations in antibody availability, we sequentially stained sections with anti-Nopp140 and anti-Npm1 antibodies to validate the existence of the residual CBs. In WT neurons, discrete Nopp140-positive but Npm1-negative foci, indicative of genuine CBs, were observed (arrows in **Figure 1E, F**). However, in *Coil^7aa^*mutant neurons, all Nopp140-positive foci were also positive for Npm1, indicating that these structures were nucleoli rather than residual CBs (**Figure 1E, F**). Taken together, these results confirm that our newly generated *Coil^7aa^*/*Coil^7aa^*mice represent a complete functional knockout that fails to form both canonical and residual Cajal bodies.

**Figure 1.**
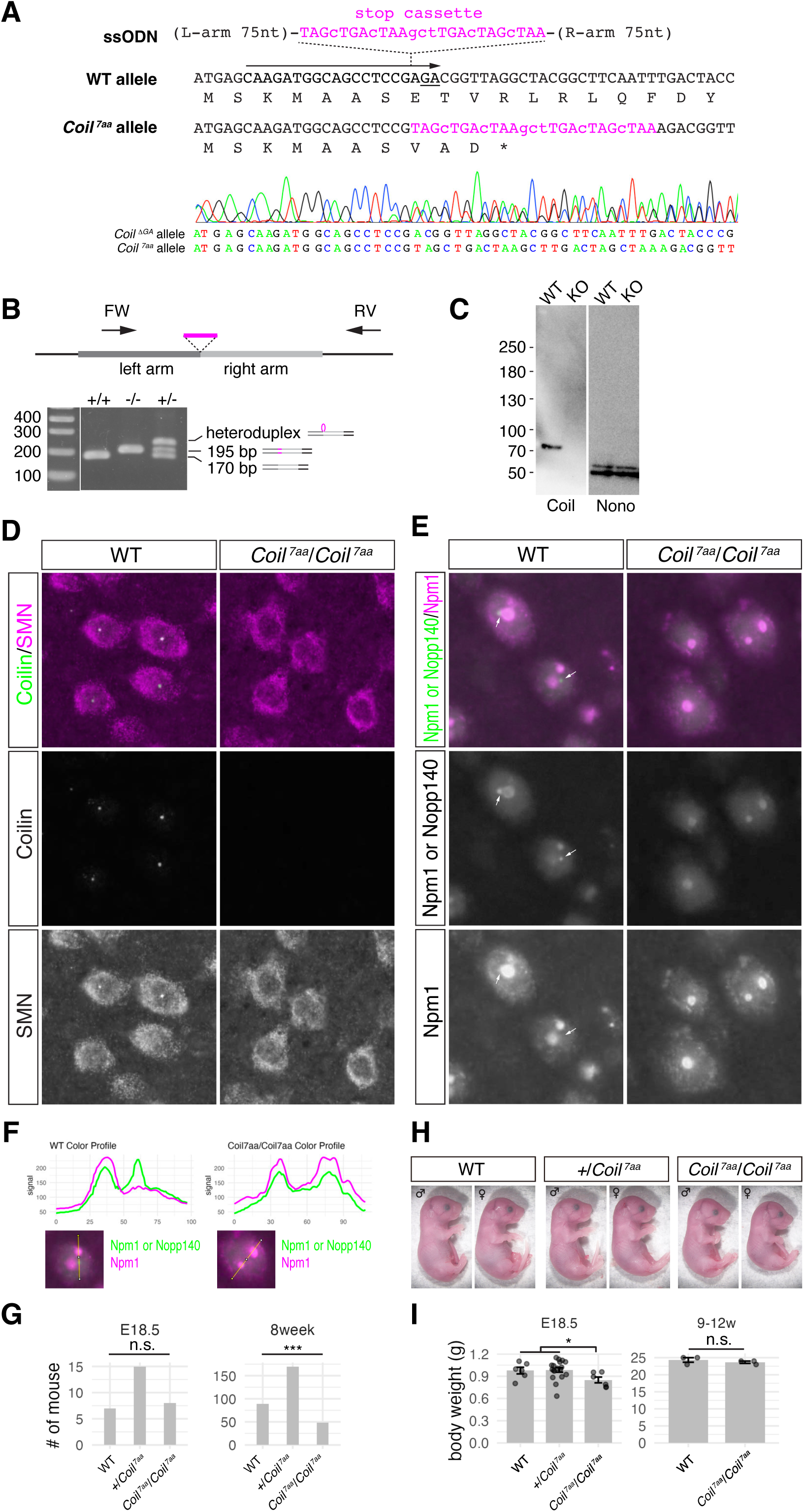
Generation of *Coil^7aa^*/*Coil^7aa^*mice and loss of Cajal body formation in mutant mice. (A) Schematic illustration of the generation of *Coil^7aa^*/*Coil^7aa^*mice by genome editing, accompanied by representative Sanger sequencing results. The designed allele includes an inserted stop cassette, and the insertion-deletion allele (*Coil^ΔGA^*) lacks two nucleotides (GA, underlined). (B) Schematic indicating primer positions used for PCR genotyping of *Coil^7aa^* mice and representative agarose gel electrophoresis patterns of amplified products. The schematic on the right illustrates how mismatched wild-type and mutant strands form a heteroduplex with a looping-out structure, leading to retarded migration during electrophoresis. (C) Immunoblot analysis confirming the absence of coilin expression in *Coil^7aa^*/*Coil^7aa^*mouse brain lysates. (D) Immunofluorescent staining of coilin (green) and SMN (magenta) in cortical neurons from WT and *Coil^7aa^*/*Coil^7aa^*mice. Note the absence of SMN-positive CBs in mutant neurons. (E) Immunofluorescent staining for Npm1 and Nopp140 in cortical neurons from WT and *Coil^7aa^*/*Coil^7aa^* mice. Due to the primary antibodies for both proteins being derived from mice, sequential staining was performed to exclusively visualize Npm1 (magenta) and subsequently both Npm1 and Nopp140 (green). Note the absence of Nopp140-only signals in *Coil^7aa^*/*Coil^7aa^* neurons, suggesting that residual CBs do not form in the mutant mice. Arrows indicate Nopp140 only signals in CBs of WT. (F) Fluorescence intensity profile plots illustrating the loss of Nopp140 signals that do not overlap with Npm1 signals in *Coil^7aa^*/*Coil^7aa^*mice. Such signals indicate CBs or residual CBs. (G) Mendelian ratios of WT, heterozygous (*+*/*Coil^7aa^*), and homozygous (*Coil^7aa^*/*Coil^7aa^*) mutant embryos at E18.5 and adults at 8 weeks. Note the significant reduction in homozygous mutant numbers at 8 weeks (***P<0.001, Chi-square test), whereas this difference was not detected at E18.5 (n.s., not significant). (H) Gross morphology of E18.5 embryos. Note that no overt abnormalities were observed in *Coil^7aa^*/*Coil^7aa^* embryos, although they were slightly smaller compared to WT or heterozygous littermates. (I) Comparison of body weight among WT, heterozygous, and *Coil^7aa^*/*Coil^7aa^*embryos at E18.5 and adult mice at 9-12 weeks. Reduced body weight was observed only at the embryonic stage, and *Coil^7aa^*/*Coil^7aa^*mice that survived to adulthood had body weights comparable to WT littermates.

### Coilin-deficient mice exhibit partial perinatal lethality but normal adult survival

Previous studies have demonstrated that coilin knockout mice exhibit a semi-lethal phenotype, with partial lethality occurring between E13.5 and P1. Consequently, homozygous mutants were recovered at approximately half the expected Mendelian ratio (Tucker et al. 2001; Walker et al. 2009). Consistent with these findings, we observed that the proportion of surviving *Coil^7aa^*/*Coil^7aa^* mice at 8 weeks of age was reduced to 54% of that of WT littermates (**Figure 1G**). To more precisely determine the developmental period during which lethality occurs, we examined embryos at E18.5. At this stage, we recovered *Coil^7aa^*/*Coil^7aa^* mutant embryos at normal Mendelian ratios comparable to those of WT littermates, with no overt morphological abnormalities (**Figure 1G, H**). However, mutant embryos were slightly smaller than their WT or heterozygous littermates, a size difference not apparent in adult mice (**Figure 1I**). These observations suggest that a subset of *Coil^7aa^*/*Coil^7aa^*mice die at birth or shortly thereafter, whereas survivors are indistinguishable from WT or heterozygous littermates once they reach adulthood, as previously reported (Tucker et al. 2001; Walker et al. 2009).

### Coilin is essential for maintaining efficient 2′-O-methylation of Sm-snRNAs *in vivo*

CBs have long been implicated in the 2′-*O*-methylation and pseudouridylation of Sm-associated snRNAs. However, previous studies using coilin mutant animal models have shown that modifications of Sm-associated snRNAs can still occur in coilin-deficient *Drosophila melanogaster* (Deryusheva and Gall 2009) and in mouse embryonic fibroblasts derived from coilin-mutant mice (Jady et al. 2003). These earlier studies indirectly assessed snRNA modifications using primer extension assays. To directly and quantitatively evaluate snRNA modifications, we performed liquid chromatography coupled with mass spectrometry (LC-MS), an approach previously optimized for the precise detection of modified nucleotides in tRNAs and other RNAs (Suzuki et al. 2007). Because the quantitative LC-MS analysis of purified snRNAs requires a substantial amount of starting material, we utilized liver tissue, which practically provides a sufficient RNA yield necessary for this assay. We isolated all spliceosomal snRNAs of the major (GU-AG) pathway—U1, U2, U4, U5, and U6—from total RNAs prepared from the livers of WT and *Coil^7aa^*/*Coil^7aa^* mice, and digested them with RNase T_1_ and/or RNase A (**Figure 2A**). The resulting RNA fragments, with or without 2′-*O*-methyl modifications, were quantified based on peak intensity in LC-MS analyses. For U2 snRNA, we focused on fragments containing Cm61, Um47, Cm40, Gm25, and m^6^Am30 (**Supplemental Fig. S1A**). Notably, methylation at Cm61 was substantially reduced in *Coil^7aa^*/*Coil^7aa^* mice compared to WT controls (82% modified in WT versus 18% in mutants; **Fig. 2B**). Similar reductions were observed at Um47, Cm40, Gm25 and m^6^Am30, where nearly all fragments were modified in WT mice, whereas 32%, 35%, 26%, and 37%, respectively, remained unmodified or hypomodified as m^6^A30 in *Coil^7aa^*/*Coil^7aa^*mice (**Fig. 2B**). Reduced 2′-*O*-methylations were also observed in U1 and U4 snRNAs, where the unmodified U1 Am70 fragment increased from 6.7% to 29.6% and the unmodified U4 Am65 increased from 0.59% to 2.3% in *Coil^7aa^*/*Coil^7aa^*mice (**Supplemental Fig. S1B-D**). In contrast, modifications at the first and second nucleotide positions (Am1 and Um2), introduced by enzymes that are not enriched in CBs (Werner et al. 2011), were unaffected (**Fig. 2B**). Likewise, modification of U6 snRNA (**Supplemental Fig. 1E**)—which is processed through a distinct nucleolar pathway independent of Sm-snRNAs—was also unaffected (**Fig. 2C)**, consistent with previous findings suggesting that U6 modification occurs predominantly in nucleoli rather than in CBs (Ganot et al. 1999; Darzacq et al. 2002). To rigorously evaluate the changes in snRNA modification levels upon *Coil* deletion, we performed comprehensive statistical analysis of these modifications using a different set of animals (**Supplemental Fig. S2**). Because only a single wild-type (WT) replicate was available for the batch of animals used in this analysis, heterozygous (*Coil^7aa/+^*) mice were included in the control group. The heterozygotes retain one functional coilin allele, and the single WT sample showed snRNA modification levels comparable to those of the heterozygous samples at every site examined, while all control samples were clearly distinct from *Coil^7aa^*/*Coil^7aa^* mice at the affected sites (**Fig. 2D**; **Supplemental Table 1**). We therefore treated WT and heterozygous animals together as a single coilin-proficient control group ("WT/Hetero," n = 3), rather than performing an underpowered statistical comparison between a single WT and the heterozygous samples. Using a two-tailed Welch’s t-test, we confirmed significant reductions in specific modification fractions in *Coil^7aa^*/*Coil^7aa^*mice compared to WT/Hetero controls (**Fig. 2D**). These findings firmly establish that coilin is selectively required for the maintenance of specific snRNA modifications *in vivo*, demonstrating that the observed phenotypes are driven by robust and statistically significant biochemical alterations rather than random biological fluctuation.

**Figure 2.**
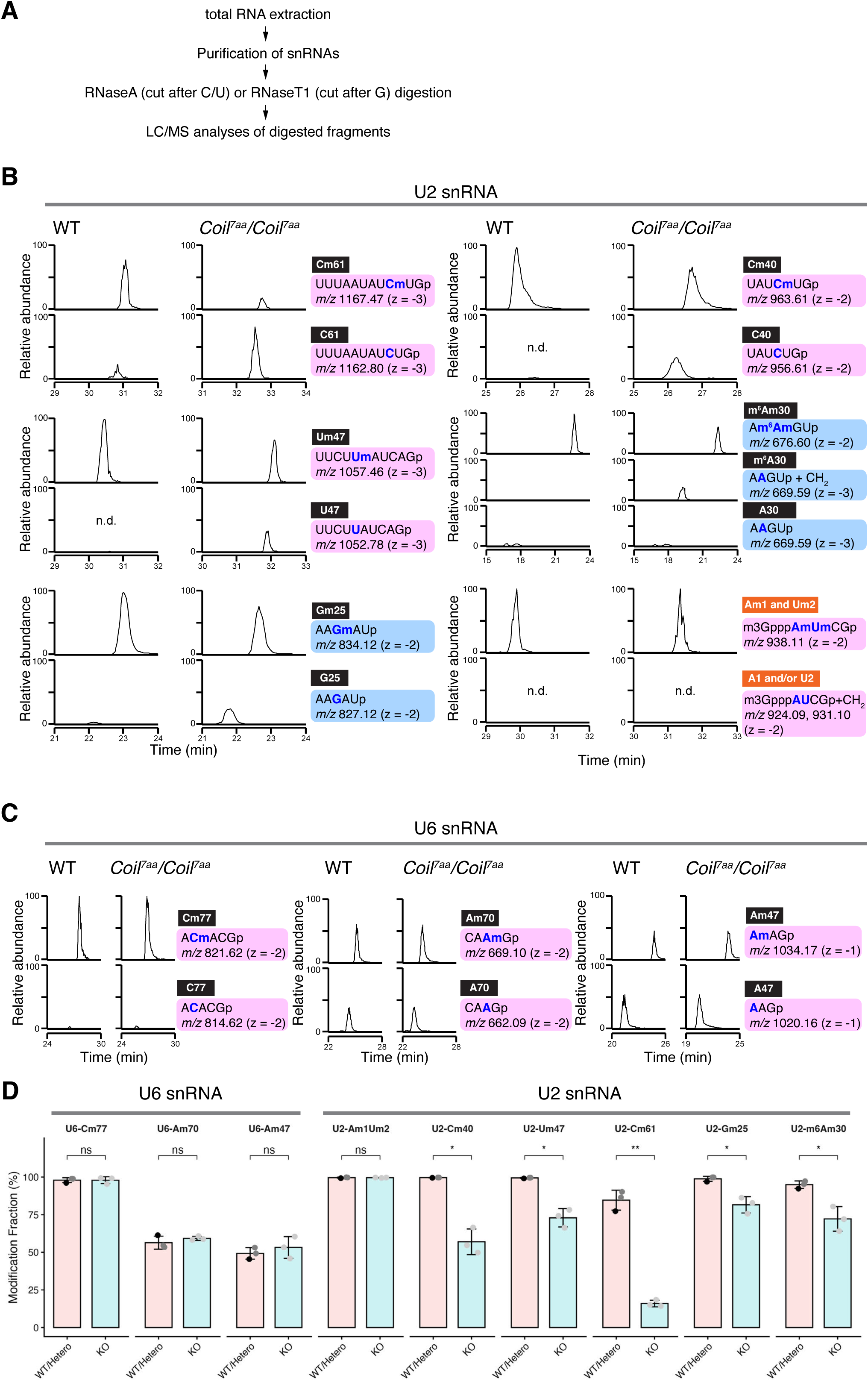
Modification of U2 snRNA is reduced in *Coil^7aa^*/*Coil^7aa^* mouse liver. (A) Schematic representation of the snRNA modification analysis workflow. Total RNA is extracted and subjected to the purification of specific snRNAs. The purified snRNAs are then enzymatically digested using RNase A (which specifically cleaves the phosphodiester bond on the 3’ side of pyrimidine residues, C and U) or RNase T1 (which specifically cleaves on the 3’ side of guanine residues, G). The resulting sequence-specific RNA fragments are subsequently analyzed by LC-MS to identify and quantify RNA modifications. (B) Extracted-ion chromatograms of RNase T_1_- and A- treated U2 snRNA fragments containing modifications Cm61, Cm40, Um47, m^6^Am30, Gm25 and Am1/Um2. Note that in all shown positions except the cap Am1/Um2 nucleotides, fragments containing unmodified nucleotides increased in *Coil^7aa^*/*Coil^7aa^*mice, accompanied by a reduction of fragments with modifications. n.d., not detected. (C) Extracted-ion chromatograms of U6 snRNA fragments containing modifications at Cm57, Am70, and Am47. The ratios of modified versus unmodified fragments remained unchanged between wild-type and *Coil^7aa^*/*Coil^7aa^* mice. (D) Quantification of snRNA modifications in *Coil^7aa^*/*Coil^7aa^* mice. Data are presented as mean ± SD from three independent biological replicates (*n* = 3). Each animal is plotted as an individual data point, with the single wild-type sample in black, the two heterozygous samples in dark gray, and the three *Coil^7aa^/Coil^7aa^*samples in light gray (n = 3 control: 2 heterozygous + 1 WT; n = 3 *Coil^7aa^/Coil^7aa^*). Statistical significance between WT/Hetero and KO groups was determined using a two-tailed Welch’s t-test. * and ** indicate *p*-values < 0.05 and 0.01, respectively; ns, not significant.

### Pseudouridylation and pre-mRNA splicing are largely insensitive to coilin depletion

While 2′-*O*-methylation is guided by C/D box scaRNAs, pseudouridylation is guided by H/ACA scaRNAs and catalyzed by dyskerin (Ganot et al. 1997; Lafontaine et al. 1998; Huttenhofer et al. 2001). To investigate the role of CBs on pseudouridylation, we employed 2-bromoacrylamide (2-BAA)-assisted cyclization sequencing (BACS), which quantifies modification levels by calculating the ratio of U-to-C conversions upon 2-BAA treatment (Xu et al. 2024). In contrast to the clear reduction in 2′-*O*-methylation, pseudouridylation levels were largely unchanged in *Coil^7aa^*/*Coil^7aa^*mice, and the few residues showing statistically significant differences did not follow a uniform trend: pseudouridylation at U60 of U2 snRNA was increased, whereas that at U53 was decreased in *Coil^7aa^*/*Coil^7aa^* mice, respectively (**Supplemental Fig. S3A, B**). These observations are consistent with a previous report showing that pseudouridylation is only modestly affected by loss of CB-localized guide scaRNAs upon Nopp140 depletion (Bizarro et al. 2021). Together, these findings suggest that H/ACA scaRNP–mediated pseudouridylation is less dependent on intact Cajal bodies than C/D box scaRNP–mediated 2′-*O*-methylation. The heterogeneous changes in pseudouridylation in *Coil^7aa^*/*Coil^7aa^* mice might arise from site-specific differences in H/ACA scaRNP dependence on Cajal bodies and/or altered competition with C/D box scaRNPs after CB disassembly.

To examine whether the altered snRNA modifications observed in *Coil^7aa^*/*Coil^7aa^*mice affected tissue histology, we analyzed hematoxylin and eosin-stained paraffin sections from WT and *Coil^7aa^*/*Coil^7aa^* livers. All expected cell types were present, and we observed no overt abnormalities in cellular organization or histological architecture (**Fig. 3A**). Next, we performed RNA sequencing (RNA-seq) analyses on liver samples from WT and *Coil^7aa^*/*Coil^7aa^*mice to investigate potential gene expression changes resulting from coilin depletion. Only 30 RefSeq-annotated genes exhibited statistically significant differential expression (padj < 0.01, log2FoldChange > 1), indicating minimal global effects on gene expression (**Fig. 3B, Supplemental Table 2**). We noticed that some of these differentially expressed genes belong to specific gene families, namely the cytochrome P450 4a (Cyp4a) and Major urinary proteins (Mup) families, which are clustered on the proximal and distal regions of chromosome 4, respectively (**Fig. 3C**). Since snRNAs are generally involved in regulating pre-mRNA splicing, we assessed splicing patterns in liver and testis using two complementary, methodologically independent tools: rMATS, which detects alternative splicing from splice-junction reads (Shen et al. 2014), and SUPPA2, which infers percent-spliced-in (PSI) values from transcript-level quantification (Trincado et al. 2018). We analyzed RNA-Seq data obtained from testis in addition to liver for the analyses because highest expression of coilin is observed in testis (Tucker et al. 2000). Differential splicing between *Coil^7aa^*/*Coil^7aa^* and control mice was evaluated separately for each tissue using replicate-based statistics (*n* = 3 per genotype). At matched thresholds (rMATS FDR < 0.01, SUPPA2 p < 0.01, |ΔPSI| ≥ 0.1), rMATS identified 73 significant genes in liver and 251 in testis, while SUPPA2 identified 4 and 8, respectively (Fig. S4A, B). Strikingly, the two tools shared no significant genes in either tissue (0 of 4,542 testable genes in liver and 0 of 7,267 testable genes in testis; Fig. S4C). Because low between-tool concordance is a recognized hallmark of false-positive–dominated splicing calls (Olofsson et al. 2023), the complete absence of events reproducibly detected by two independent methods indicates that no robust, coherent splicing program is altered in these tissues. These observations suggested that the reduction in snRNA modifications observed in *Coil^7aa^*/*Coil^7aa^*mice does not substantially alter pre-mRNA splicing patterns in the liver and testis examined here under standard laboratory conditions. Because our RNA-seq analysis was restricted to these two tissues, we cannot exclude tissue- or condition-specific splicing effects in other tissues, which remain to be addressed in future studies. Collectively, we conclude that despite the reduction in specific 2′-O-methylations, the absence of coilin does not substantially alter global pseudouridylation or functionally compromise pre-mRNA splicing fidelity under standard physiological conditions.

**Figure 3.**
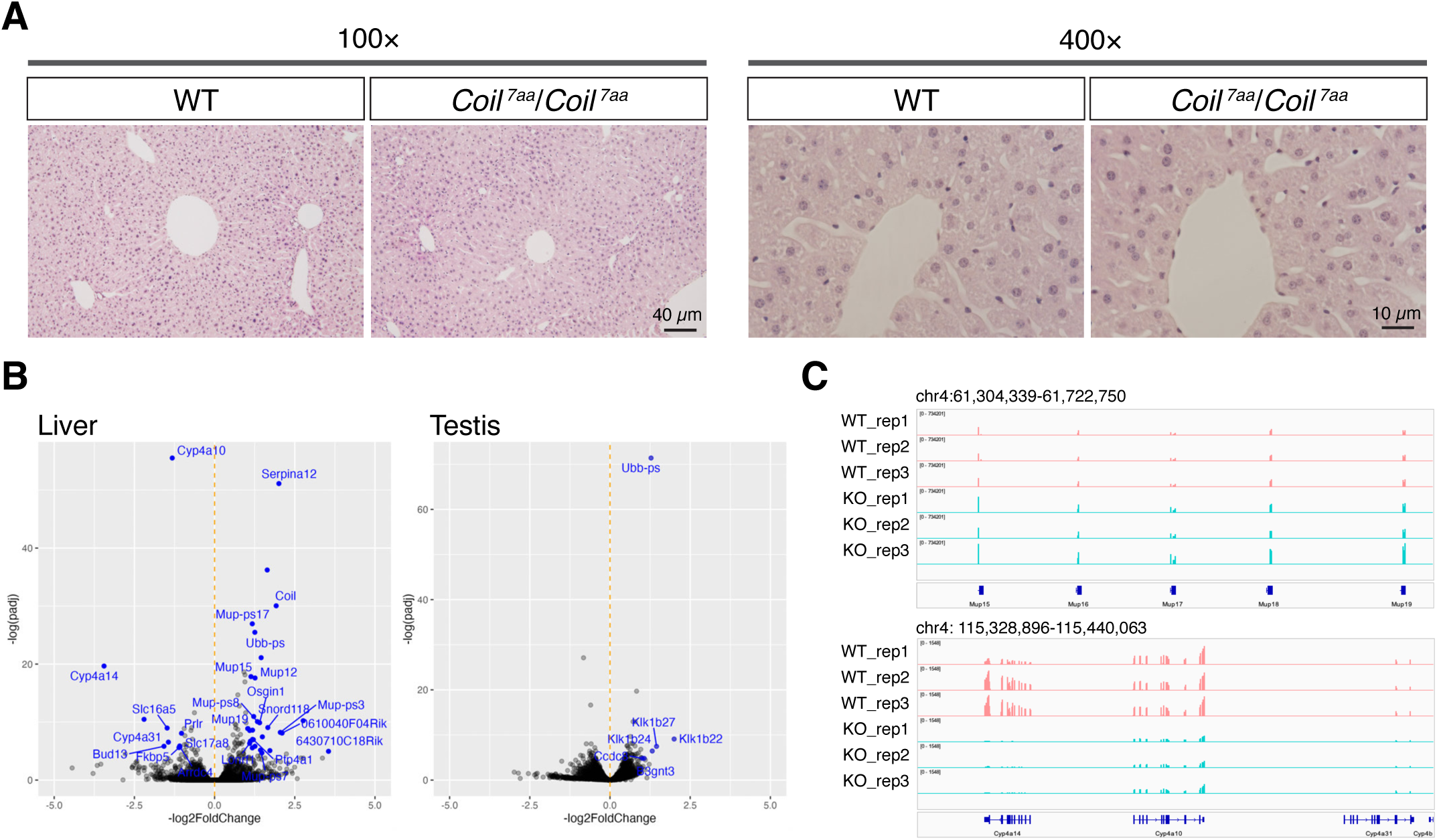
Histology and gene expression are minimally affected in *Coil^7aa^*/*Coil^7aa^* mouse liver. (A) Representative images of hematoxylin-eosin (HE)-stained histological sections from wild-type (WT) and *Coil^7aa^*/*Coil^7aa^* mouse livers at low (100×) and high (400×) magnifications. No gross abnormalities were observed. (B) Volcano plots showing differential gene expression between WT and *Coil^7aa^*/*Coil^7aa^* mouse liver and testis. Only a small number of genes were significantly altered. Blue dots indicate significantly differentially expressed genes (padj < 0.01, log2FoldChange > 1). (C) Genome browser views of RNA-seq reads mapped to regions where differentially expressed genes form clusters. Note the increased expression of the Mup and Klk gene families and decreased expression of the Cyp4a gene family in *Coil^7aa^*/*Coil^7aa^*mice.

### High-quality sperms are reduced in *Coil^7aa^*/*Coil^7aa^* mice

Previous studies have shown that *Coil^82aa^*/*Coil^82aa^*mice exhibit reduced fertility, despite normal histological appearances of testes and ovaries (Walker et al. 2009). Given that coilin expression is highest in the testis (Tucker et al. 2000), we revisited this tissue in more detail in our *Coil^7aa^*/*Coil^7aa^* mice. Consistent with previous observations, testis size in *Coil^7aa^*/*Coil^7aa^* mice was smaller compared to WT littermates (**Fig. 4A, B**). To investigate this further, we examined histological sections of seminiferous tubules at various stages of spermatogenesis. All expected cell types, including spermatogonia, spermatocytes, round spermatids, and elongated spermatids, were normally arranged without noticeable abnormalities (**Fig. 4C**). RNA-seq analyses revealed minimal overall changes in gene expression; however, some differentially expressed genes belonged to the kallikrein (Klk) gene family (**Fig. 3B**), which forms a gene cluster on chromosome 7. We also examined the gross morphology of mature sperm collected from the epididymis of *Coil^7aa^*/*Coil^7aa^* mice, and observed no morphological defects by light microscopy (**Fig. 4D**).

**Figure 4.**
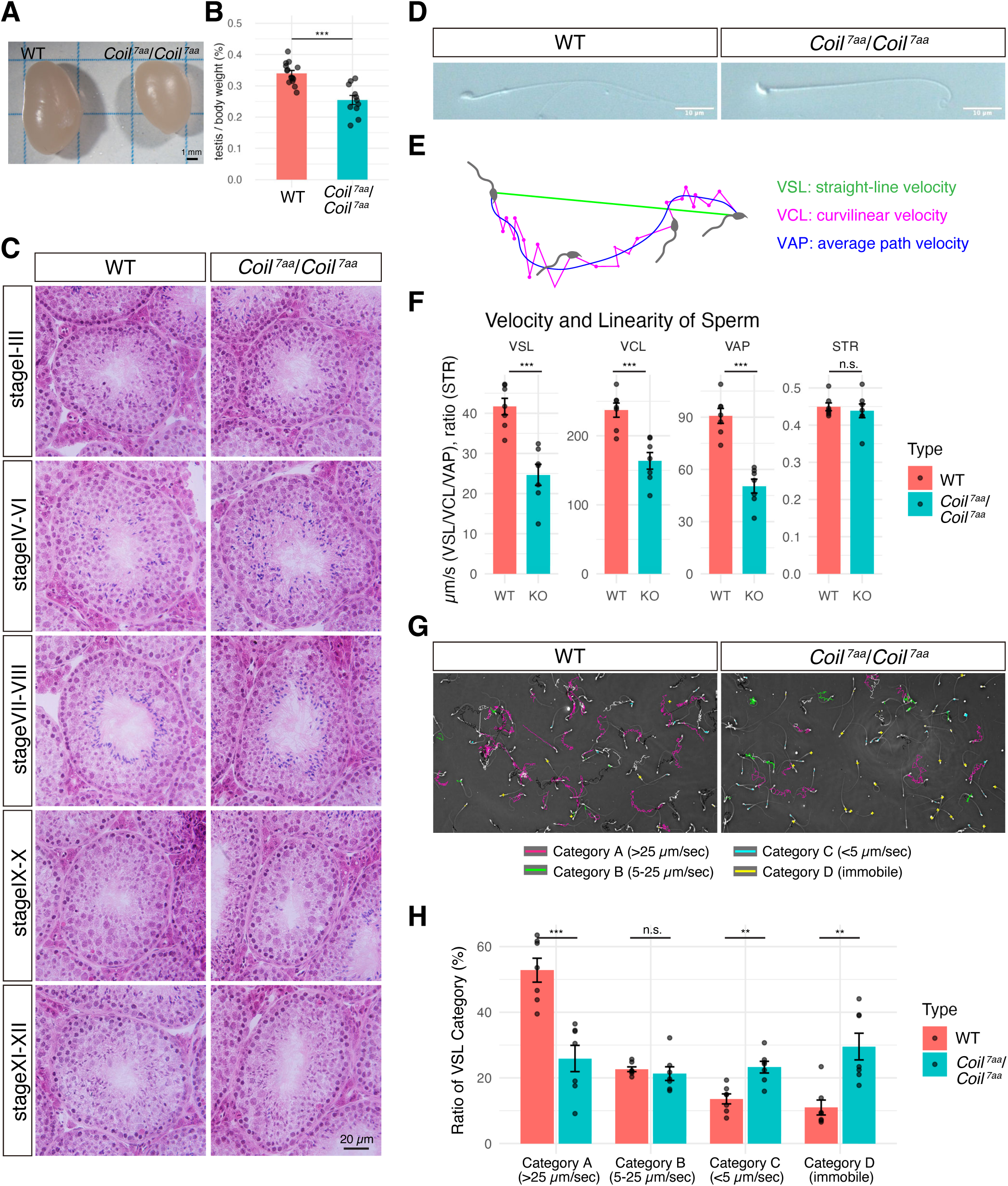
Reduced sperm motility in *Coil^7aa^*/*Coil^7aa^* male mice compared to wild-type mice. (A) External appearance of testes dissected from wild-type (WT) and *Coil^7aa^*/*Coil^7aa^*mice. (B) Quantification of testis weight normalized by body weight. Note the significant reduction in *Coil^7aa^*/*Coil^7aa^* mice. (C) HE-stained histological sections of seminiferous epithelia from WT and *Coil^7aa^*/*Coil^7aa^*mice at various stages of spermatogenesis. No observable abnormalities were detected at this level. (D) Differential interference contrast (DIC) microscopy images of mature sperm obtained from the epididymides of WT and *Coil^7aa^*/*Coil^7aa^*mice. Gross morphological defects were not evident. (E) Schematic illustrating the measurement of sperm velocities. Three parameters are shown: VSL (straight-line velocity), which measures the linear distance between the initial and final positions of sperm heads during the observation period; VCL (curvilinear velocity), measuring the total distance traversed along the sperm’s actual trajectory; and VAP (average path velocity), measuring the distance traveled along a smoothed, average path. (F) Quantification of sperm velocity in WT and *Coil^7aa^*/*Coil^7aa^* mice using the three velocity parameters. Note the decreased velocity across all measurements in *Coil^7aa^*/*Coil^7aa^* sperm compared to WT. However, the straightness (STR), calculated as VSL divided by VAP, was not significantly different. (G) Representative sperm movement trajectories from WT and *Coil^7aa^*/*Coil^7aa^* mice, color-coded according to sperm speed: category A (>25 µm/sec, magenta), category B (5–25 µm/sec, white), category C (<5 µm/sec, green), and category D (immotile, yellow cross). (H) Quantification of sperm motility categories. Each dot represents an individual mouse, with error bars indicating standard errors. The proportion of active sperm (category A) was significantly reduced, whereas the immotile fraction (category D) was increased in *Coil^7aa^*/*Coil^7aa^*mice (**P<0.01, ***P<0.001, Wilcoxon rank-sum test).

We then assessed sperm motility using a sperm motility analysis system (SMAS). We quantitatively analyzed sperm motility based on several velocity parameters: straight-line velocity (VSL), representing the straight-line distance traveled by the sperm per unit time; curvilinear velocity (VCL), measuring the actual trajectory speed; and average path velocity (VAP), calculated along a smoothed trajectory. Notably, *Coil^7aa^*/*Coil^7aa^* sperm showed significant reductions in all three parameters compared to WT sperm (**Fig. 4E-H**). In contrast, straightness (STR)—the ratio of VSL to VAP—was not significantly different, indicating reduced sperm speed without substantial changes in trajectory linearity (**Fig. 4F**). To further characterize sperm motility, we categorized sperm into four groups based on VSL values: rate A (>25 µm/sec), rate B (5–25 µm/sec), rate C (<5 µm/sec), and rate D (immotile) (**Fig. 4G**). In WT mice, 52.8% of sperm belonged to the most active group (rate A), while this proportion markedly decreased to 25.9% in *Coil^7aa^*/*Coil^7aa^*mice (**Fig. 4H**). Conversely, the immotile sperm group (rate D) increased significantly in *Coil^7aa^*/*Coil^7aa^*mice (29.5% compared to 11.0% in WT) (**Fig. 4H**). Although our preliminary mating test showed that *Coil^7aa^*/*Coil^7aa^* males are not completely sterile and are capable of producing offspring, we observed substantial variance in litter sizes among the limited number of replicates (n = 3; litter sizes: wildtype, 6.6 ± 1.5; *Coil^7aa^*/*Coil^7aa^*, 6.0 ± 2.6). A previous study using a different coilin mutant model (*Coil^82aa^*/*Coil^82aa^*) reported a partial infertility phenotype, describing the mice as "reproductively less fit" with reduced litter sizes when evaluated over a larger cohort and a longer duration (6 months; Walker et al., 2009). Given the small sample size and short duration of our current assessment, our data are insufficient to quantitatively exclude the possibility of a similar subtle decline in reproductive fitness. Therefore, whether the clean deletion of the *Coil* causes partial infertility remains an open question that requires future large-scale and long-term mating assessments. These results indicate that coilin is necessary for producing high-quality sperm, a factor potentially underlying the reduced fertility phenotype previously observed in *Coil^82aa^*/*Coil^82aa^* mice.

### Cajal body formation is restricted to a subset of cells during spermatogenesis

To investigate the molecular mechanisms underlying the decreased sperm motility observed in *Coil^7aa^*/*Coil^7aa^* mice, we examined the expression patterns of coilin during spermatogenesis. For this purpose, testis sections were prepared from tissues freshly frozen in isopentane chilled in liquid nitrogen. Rapid freezing was essential to preserve antigenicity of protein markers and histological integrity, facilitating accurate cell-type identification. Specific stages of seminiferous tubules were identified based on the characteristic staining patterns of peanut agglutinin (PNA) (Nakata et al. 2015) and gamma histone H2AX (γH2AX) (Blanco-Rodriguez 2009) and coilin (**Fig. 5A**), as detailed in the Materials and Methods and **Supplemental Table 3**.

**Figure 5.**
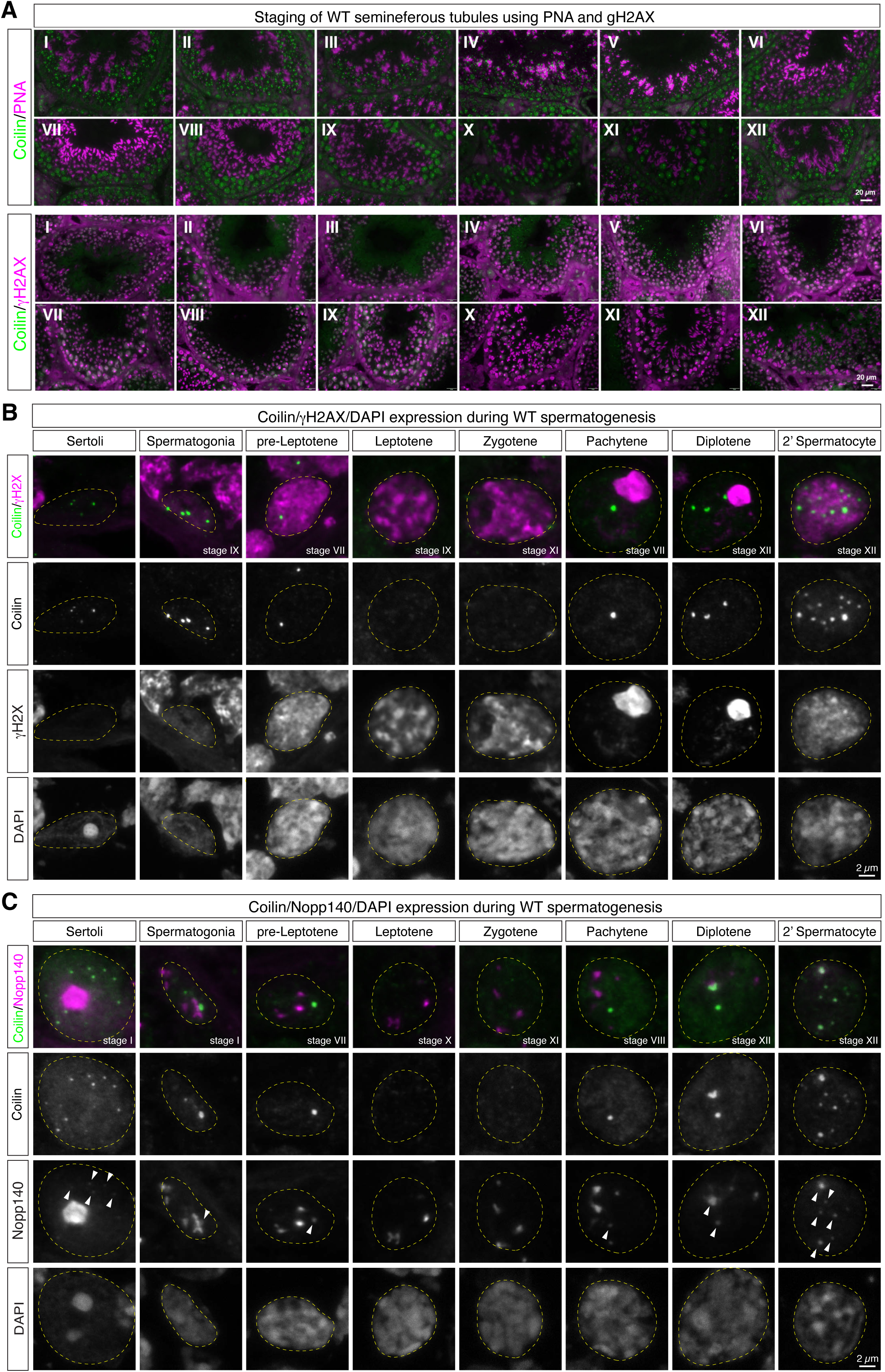
Formation of CBs during mouse spermatogenesis. (A) Representative images showing seminiferous tubule stages identified by distinct subcellular localization patterns of PNA and γH2AX staining. Stages (I–XII) were precisely determined by combined staining patterns of PNA, γH2AX, DAPI, and coilin. (B) Representative images illustrating the dynamic expression patterns of coilin (green), γH2AX (red), and DAPI (blue) across different seminiferous tubule stages. Note the absence of coilin expression in leptotene and zygotene spermatocytes, with expression reappearing at the pachytene stage. (C) Identification of CBs based on the co-localization of coilin (green) with Nopp140 (red) in seminiferous tubules during spermatogenesis. Nopp140-positive nucleoli lacking coilin signals gradually diminish and eventually disappear during later spermatocyte stages (diplotene and secondary spermatocytes), consistent with known nucleolar fragmentation at these stages. Arrowheads indicate positions of Cajal bodies identified by co-localization of coilin and Nopp140. Yellow dotted lines indicate the position of nuclei.

Sertoli cells were distinguished by their pale nuclei, and displayed multiple small round CBs characterized by the colocalization of coilin and Nopp140, the latter also marking nucleoli (**Fig. 5B, C**). Spermatogonia were identified by their oval nuclei, lack of γH2AX staining, and location along the basement membrane in the outermost layer of the seminiferous epithelium. Within this cell type, coilin-positive CBs were detected in a small subset of cells (1-2 spermatogonia/seminiferous tubule), typically appearing as 1-3 foci per nucleus, larger than those observed in Sertoli cells and other somatic cell types (**Fig. 5B, C**). As spermatogonia commit to meiosis and differentiate into pre-leptotene spermatocytes, they move toward the adluminal compartment and begin expressing γH2AX. Coilin expression was maintained in pre-leptotene spermatocytes at stages VII/VIII but disappeared during the subsequent leptotene and zygotene stages (**Fig. 5B, C**). Prominent coilin expression resumed in pachytene spermatocytes, characterized by a γH2AX-positive XY body, and persisted throughout the pachytene stage. In these cells, coilin formed distinct single round CBs overlapping with Nopp140 (**Fig. 5C**). Expression of coilin was further elevated in diplotene spermatocytes, specifically at stage XI, where an increased number of coilin-positive CBs was observed, accompanied by a decrease in nucleoli defined by Nopp140 signals without coilin (**Fig. 5C**). In secondary spermatocytes, present specifically at stage XII, coilin expression was even further elevated, forming multiple CBs that overlapped precisely with Nopp140 (**Fig. 5C**). At this stage and beyond, nucleoli identified by Nopp140 staining without coilin signals disappeared, consistent with a previous report describing nucleolar fragmentation and disappearance during late meiosis stages (Czaker 1985; Biggiogera et al. 1991). These observations demonstrate that coilin expression and Cajal body formation are highly dynamic and strictly regulated throughout spermatogenesis, exhibiting a transient disappearance during early meiosis followed by prominent upregulation in late meiotic spermatocytes.

### Coilin is transiently localized to the nuclear pocket in elongating spermatids

Later stages of spermatogenesis, where round spermatids differentiate into elongated sperm, are collectively referred to as spermiogenesis, and categorized into steps 1 to 16 (Russell et al. 1990), which stages are clearly distinguishable by characteristic PNA staining patterns (Nakata et al. 2015) (**Supplemental Table 3**). During these stages, coilin also exhibited a dynamic expression pattern (**Fig. 6A**). In round spermatids characterized by a centrally located, single, round heterochromatic structure stained by DAPI, coilin formed 2–4 distinct bright foci overlapping with Nopp140 signals (**Fig. 6B**). As spermatids differentiated into elongated forms, these nuclear CBs disappeared. The expression re-started at later steps, and coilin transiently formed distinct condensates at the posterior region of the nucleus, specifically at step 15 (**Fig. 6A, B**). These coilin condensates, appearing at the late stage of spermiogenesis, did not overlap with Nopp140 signals, indicating they differ from typical nuclear CBs (**Fig. 6B**). These coilin signals rapidly diminished in subsequent spermiogenesis steps and were almost undetectable by step 16 (**Fig. 6B**).

**Figure 6.**
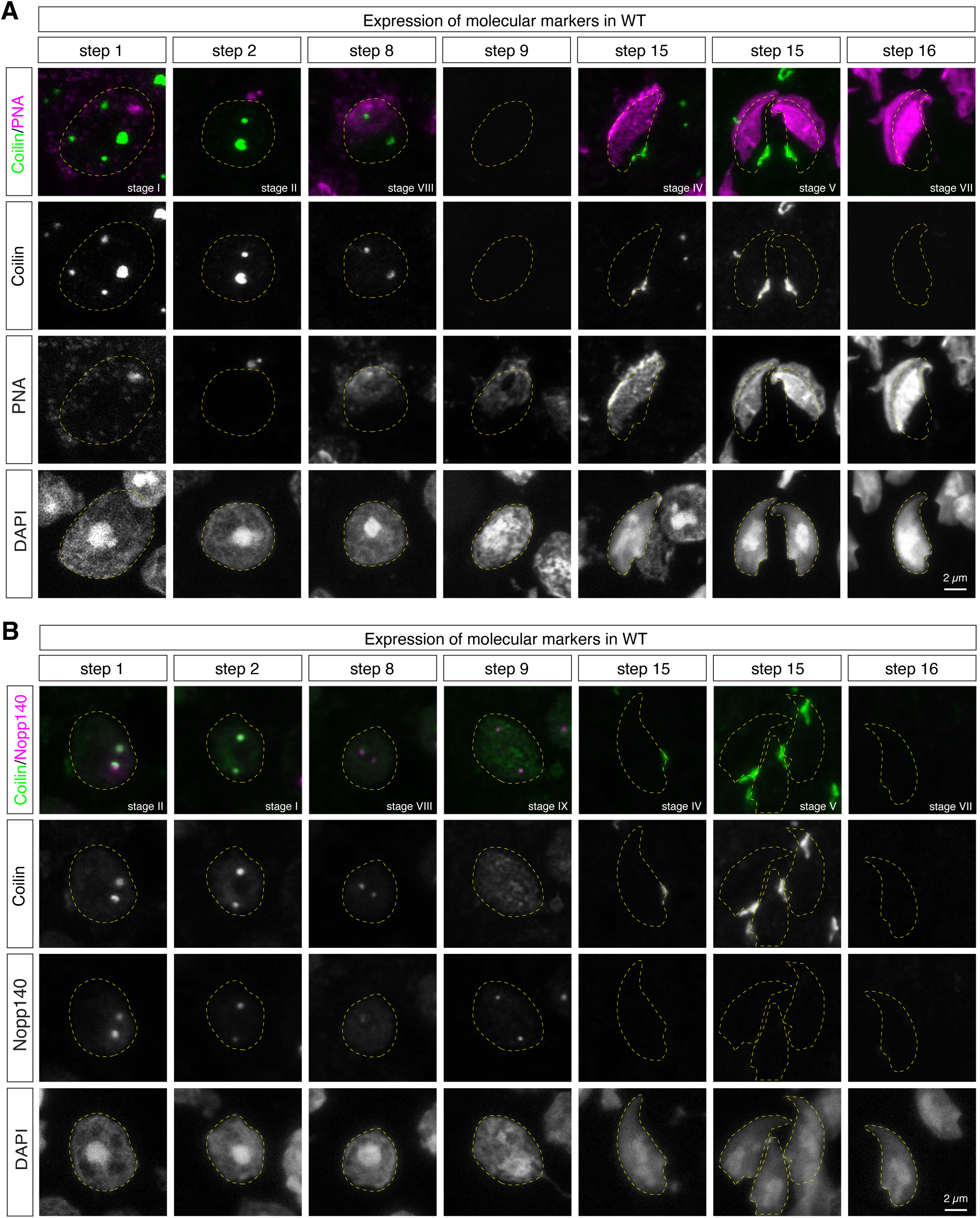
Localization of coilin during mouse spermiogenesis. (A) Representative images illustrating the dynamic localization of coilin (green), DAPI (blue), and either PNA (red in A) during the transition from round spermatids to elongating spermatids. In round spermatids, coilin initially forms multiple distinct nuclear foci overlapping with Nopp140. These nuclear signals gradually diminish and become undetectable around step 9. Coilin expression reappears around step 14, reaching maximum intensity at step 15, at which point coilin accumulates in condensates located within the nuclear pocket formed at the posterior region of elongating spermatid nuclei. Notably, these coilin-positive condensates at step 15 do not overlap with Nopp140 signals, suggesting that they are structurally and functionally distinct from canonical CBs. (B) Representative images similar to those in (A) staining for Nopp140 instead of PNA. Yellow dotted lines indicate the position of nuclei.

Because these coilin signals outside canonical CBs represented one of the most prominent expressions in seminiferous tubules (**Fig. 5A, 6A**), we further investigated their precise localization in elongating spermatids. Notably, at this stage, canonical Cajal body markers, including FBL, Nopp140, U2 snRNA, and Sca2 RNA, were completely absent in elongating spermatids (**Fig. 7A**). This indicates that the coilin-containing condensates observed in these cells are distinct from canonical Cajal bodies. During the transition from round to elongated spermatids, chromatin undergoes extensive remodeling, including histone-to-protamine replacement, resulting in highly compacted sperm nuclei (Okada 2022). Concomitantly, an electron-lucent structure called the nuclear pocket (Czaker 1985), enclosed by a redundant nuclear envelope (Lalli and Clermont 1981), transiently forms at the posterior pole of elongating spermatid nuclei around steps 14–15. Given the localization and timing of coilin expression, we hypothesized that coilin specifically localizes to this nuclear pocket. Because the redundant nuclear envelope is enriched with nuclear pore complexes (NPC) (Ho 2010; Uemura et al. 2023), we performed double-staining experiments for coilin and NPC (**Fig. 7B**). As expected, coilin signals extensively overlapped with NPC staining, confirming coilin localization within the nuclear pocket at step 15 spermatids. We also examined the distribution of Lamin B1, which has recently been reported to recede toward the posterior pole near the developing flagellum during spermiogenesis—a region closely associated with the nuclear pocket (Bragina et al. 2024). The coilin signals were detected precisely between Lamin B1 and the DAPI-positive chromatin region, further supporting our hypothesis that coilin forms condensates within the nuclear pocket (**Fig. 7B**). Importantly, localization patterns of both the redundant nuclear envelope and Lamin B1 condensates at the posterior pole of elongating spermatid nuclei were unaffected in *Coil^7aa^*/*Coil^7aa^*mice (**Fig. 7C**), suggesting that coilin is dispensable for the formation of nuclear pockets and their associated structures *per se*.

**Figure 7.**
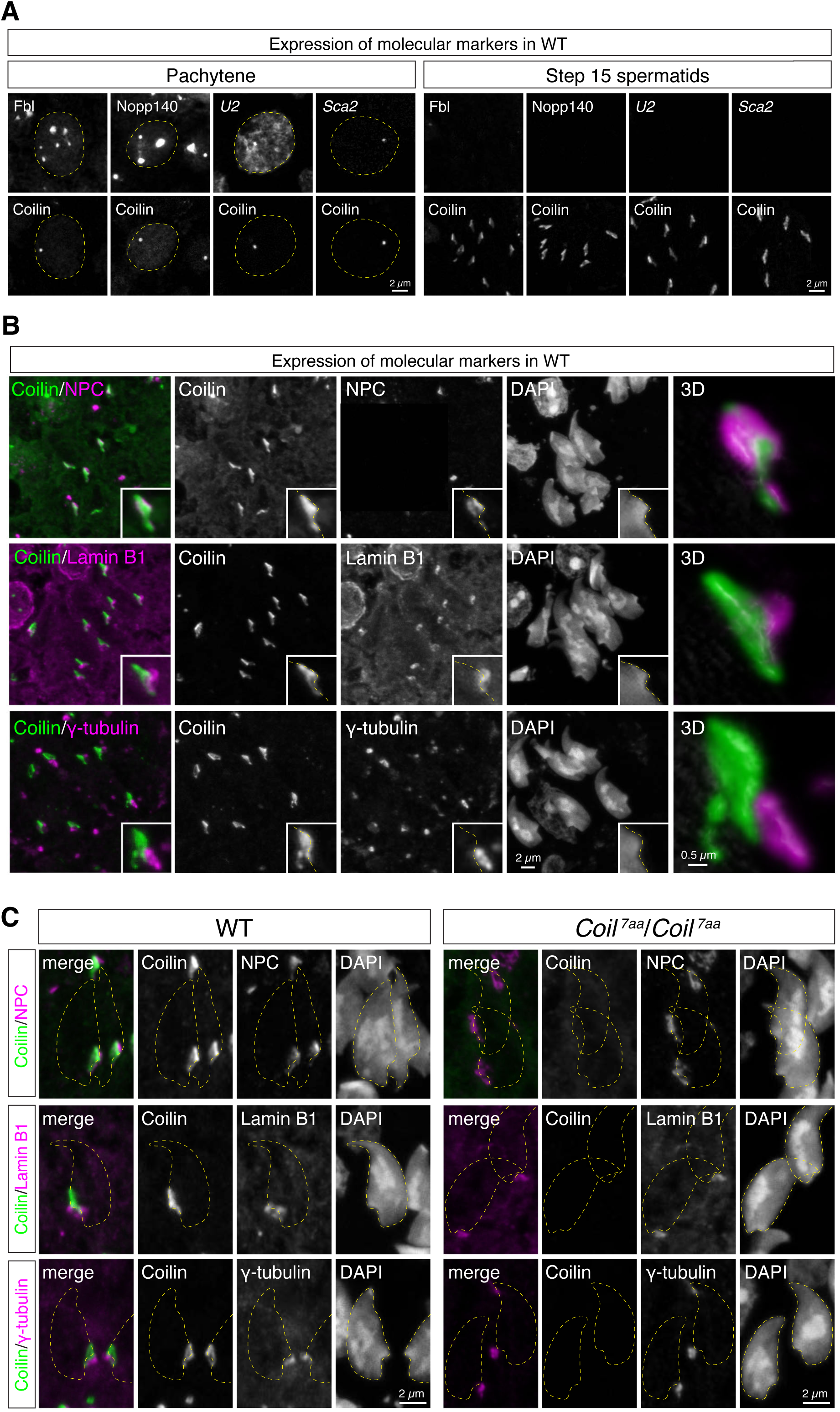
Localization of coilin to the nuclear pocket in elongating spermatids. (A) Expression and localization of canonical Cajal body markers (FBL, Nopp140, U2 snRNA, and Sca2 RNA) in pachytene spermatocytes and elongating step 15 spermatids. Note that while the signals of these markers overlapped with coilin in pachytene spermatocytes, they were undetectable in elongating spermatids harboring the coilin condensates. (B) Maximum-intensity projections and 3D rendering images (3D) of confocal microscopy images showing simultaneous detection of coilin (green), NPC/Lamin B1/γ-tubulin (red), and DAPI (blue). Coilin signals extensively overlapped with NPC staining, confirming its localization to the nuclear pocket structure surrounded by the redundant nuclear envelope. (C) Maximum-intensity projections comparing the localization of coilin (green), NPC/Lamin B1/γ-tubulin (red), and DAPI (blue) in WT and *Coil^7aa^*/*Coil^7aa^* spermatids. The positioning of nuclear pockets and centriolar complexes at the posterior pole of elongating spermatid nuclei was unaffected in *Coil^7aa^*/*Coil^7aa^* mice. Yellow dotted lines indicate the position of nuclei.

In mouse spermiogenesis, a centriolar complex containing γ-tubulin is localized adjacent to the posterior concavity of elongated spermatid nuclei (Manandhar et al. 1998), closely associated with the nuclear pocket region (Lalli and Clermont 1981; Czaker 1985). Given the observed reduction in sperm quality in *Coil^7aa^*/*Coil^7aa^* mice, we speculated that nuclear pocket-localized coilin may indirectly influence centriolar positioning and subsequent axoneme microtubule formation. To investigate this possibility, we performed simultaneous immunostaining for coilin and γ-tubulin in step-15 elongating spermatids. Although these signals were in close proximity, we did not detect direct overlap (**Fig. 7B**). Furthermore, the formation and positioning of γ-tubulin-positive centriolar complexes appeared unaffected in *Coil^7aa^*/*Coil^7aa^*mice (**Fig. 7C**). Taken together, these findings reveal a novel, non-canonical localization of coilin condensates within the nuclear pocket of late elongating spermatids, although coilin itself appears dispensable for the structural assembly of the nuclear pocket and the adjacent centriolar complexes.

## Discussion

Using quantitative LC-MS analyses, we demonstrated that coilin is required for efficient snRNA modification in mice. To our knowledge, this is the first study to explicitly confirm the functional significance of CBs in snRNA modification. Consistent with previous studies that indirectly assessed modifications using primer extension methods (Jady et al. 2003; Deryusheva and Gall 2009), we found that the amount of modified snRNA fragments was decreased but still detectable in *Coil^7aa^*/*Coil^7aa^* mice. Despite the observed reduction in snRNA modifications in the liver, we detected no overt alterations in pre-mRNA splicing patterns at least in the liver and the testis under normal laboratory conditions. It should be noted that coilin expression varies considerably among different tissues and cell types, and that the presence and size of CBs also vary. Indeed, prominent Cajal body formation accompanied by SMN condensates is limited to a small subset (∼10%) of cortical neurons, while liver cells ubiquitously exhibit CBs that are significantly smaller in size. Therefore, it is plausible that efficient snRNA modifications facilitated by coilin are particularly important for splicing in specific cell types that express coilin at high levels and form large CBs. Single-cell analyses of splicing patterns in coilin-rich neuronal populations would provide valuable insights to test this hypothesis.

Previous studies demonstrated semi-lethality in coilin knockout mice occurring between E13.5 and birth (Tucker et al. 2001; Walker et al. 2009). We further narrowed down the lethality window, demonstrating that *Coil^7aa^*/*Coil^7aa^*embryos were present at normal Mendelian ratios at E18.5, immediately before birth. Although mutant embryos exhibited slightly reduced size, no overt morphological defects were detectable anatomically at this stage. These findings suggest neonatal lethality may result from critical physiological failures occurring around birth, such as defects in initiating respiratory function or metabolic transitions due to a subtle delay in late fetal development. Further molecular and histological analyses of tissues involved in neonatal respiratory and metabolic initiation processes will be necessary to precisely determine the cause of lethality in coilin-deficient mice.

While neonatal lethality was observed in approximately half of *Coil^7aa^*/*Coil^7aa^*mutants, reduced testis size was a consistent phenotype, aligning well with high coilin expression levels observed in the testis (Tucker et al. 2000). Despite reduced testis size, histological examination revealed normal architecture and unchanged cellular composition of seminiferous tubules. Likewise, bulk RNA-seq analyses detected only a few differentially expressed genes in *Coil^7aa^*/*Coil^7aa^* testes, suggesting minimal disruption of overall gene expression. It is possible that coilin may influence the proliferation of a minor subset of cells, such as spermatogonial stem cells (Yoshida 2020), thereby reducing the total length of seminiferous tubules without altering cellular composition. Alternatively, coilin might regulate early embryonic gene programs critical for testicular development, indirectly resulting in a smaller adult testis size. Single-cell RNA sequencing studies will help distinguish these two potential mechanisms.

It is noteworthy that some of the differentially expressed genes in *Coil^7aa^*/*Coil^7aa^* mice belonged to gene families clustered at specific chromosomal loci (e.g., the Klk family on chromosome 7, and the Cyp4a and Mup families on chromosome 4). Because the effects of coilin knockout on gene expression differed between these clusters (upregulation for Klk and Mup families, downregulation for the Cyp4a family), CBs may not directly recruit these loci into transcriptionally active or repressive compartments, but instead may influence gene expression by altering chromosome organization. Consistent with this idea, a recent genome-wide chromosome conformation study demonstrated that CBs localize to multi-chromosome interfaces and mediate long-range interactions between clustered genomic regions, including highly expressed histone and U sn/snoRNA gene clusters (Wang et al. 2016). It has also been proposed that nuclear bodies formed via liquid-liquid phase separation locally modulate transcriptional activity through the concentration of transcriptional regulators or chromatin remodeling factors (Shin and Brangwynne 2017; Sabari et al. 2020). Although genomic loci associated with CBs in mice remain largely unexplored, our observations raise an intriguing hypothesis for future studies to elucidate a novel role for coilin-dependent condensates in regulating nuclear architecture and local gene expression.

The strongest coilin signals in testis were detected in the nuclear pocket of elongating spermatids at step 15 of spermiogenesis. The nuclear pocket is an electron-lucent compartment enriched with components of the ubiquitin-proteasome system, implicated in the degradation of proteins expelled from the nucleus during the histone-to-protamine transition, critical for sperm nucleus condensation (Haraguchi et al. 2007). Currently, specific nuclear substrates degraded in the nuclear pocket remain unknown, and identifying such proteins would be crucial to establish coilin’s functional role in this compartment. Notably, coilin signals appeared specifically at step 15 and were not detectable at earlier stages, indicating that the presence of coilin in the nuclear pocket is not due to degradation of residual proteins from earlier steps but reflects tightly regulated stage-specific translation. Given that translation of proteins required for later spermiogenesis stages is tightly controlled by RNA-binding proteins such as FXR1 (Kang et al. 2022), future studies should explore regulatory mechanisms governing coilin protein expression specifically at step 15, when the nuclear pocket is prominently formed.

The redundant nuclear envelope (RNE) surrounding the nuclear pocket initially forms a pouch-like structure during early spermiogenesis but subsequently becomes compressed and shrinks in later stages (Uemura et al. 2023). Thus, the abrupt disappearance of coilin signals at step 16 may result from structural alterations in the RNE. Intriguingly, the RNE has also been proposed to function as a Ca²⁺ reservoir (Uemura et al. 2023), suggesting that abnormalities in RNE integrity or function could potentially contribute to the observed decrease in sperm motility. Although no obvious changes in NPC or RNE localization were observed in coilin-deficient spermatids, subtle alterations in RNE-related functions, such as calcium signaling or protein degradation, might explain reduced sperm motility. Alternatively, recent studies have highlighted the importance of a specialized structure known as the head-tail coupling apparatus (HTCA), responsible for docking the nuclear envelope to the centrosome during spermiogenesis (Zhang et al. 2021; Wang et al. 2024). While no overt changes were seen in γ-tubulin localization or centriolar positioning in coilin knockout spermatids, it remains possible that subtle structural or functional abnormalities within the nuclear envelope or HTCA occur in the absence of coilin, potentially affecting sperm motility. Future ultrastructural and biochemical studies are needed to clarify whether such subtle alterations underlie the impaired sperm motility observed in *Coil^7aa^*/*Coil^7aa^* mice. Given the reduced sperm motility phenotype observed here, a rigorous, large-scale and long-term fertility assessment is clearly warranted to determine whether this translates into a measurable reduction in reproductive fitness; such an analysis is beyond the scope of the present study and remains an important subject for future investigation.

## Materials and Methods

### Animals

All experiments were approved by the Safety Division of Hokkaido University (Approval #2021-027) and the Animal Care and Use Committee of Hokkaido University (Approval #20-0031). *Coil^7aa^* mice were bred and maintained on a C57BL/6N genetic background under controlled laboratory conditions at 23 ± 1°C with a constant 12-hour light/dark cycle. For anesthesia, mice were intraperitoneally injected with a mixture of three anesthetics (medetomidine, midazolam, and butorphanol; 0.75, 4, and 5 mg/kg, respectively), administered at a volume of 10 µl per gram of body weight, as previously described (Kawai et al. 2011). Mice were generated either through natural mating or by in vitro fertilization followed by embryo transfer (Behringer et al. 2014).

### Genome editing

Coilin mutant mice were generated using the improved Genome editing via Oviductal Nucleic Acids Delivery (i-GONAD) method (Gurumurthy et al. 2019). Briefly, 0.3 nmol Alt-R modified crRNA (IDT) was duplexed with 0.3 nmol trRNA (IDT, #1072534) in 3 µl duplex buffer (IDT), using a thermocycler (94°C for 2 min, 85°C for 2 s, and stepwise cooling by 3 s intervals down to 25°C). The duplexed RNA was incubated with 1 µl recombinant Cas9 enzyme (IDT, #1081058) at 37°C for 10 min and then combined with 0.3 nmol Alt-R HDR-modified single-stranded oligonucleotide DNA (IDT). The assembled complex was injected into the oviduct, upstream of the ampulla, at gestational day 0.75 of ICR mice, followed by electroporation using NEPA21 (NEPA GENE) with poring pulses (50 V, 5-ms pulse, 50-ms interval, 3 pulses, 10% decay, ± pulse orientation) and transfer pulses (10 V, 50-ms pulse duration, 50-ms intervals, three pulses, and 40% decay ± pulse orientation). Obtained heterozygous *Coil^7aa^*mice were backcrossed onto a C57BL/6N genetic background for five generations prior to experiments. Sequences of crRNA and single-stranded DNA are shown in **Supplemental Table 4**.

### LC-MS analyses of snRNA modification

The small RNA fraction from mouse liver total RNA was roughly fractionated by anion exchange chromatography with DEAE Sepharose Fast Flow (GE Healthcare) (Supplementary Fig. S1A). 5′-terminal ethylcarbamate amino-modified DNA probes (**Supplemental Table 4**) covalently immobilized on NHS-activated Sepharose 4 Fast Flow (GE Healthcare) were used for snRNA isolation by RCC (Miyauchi et al. 2007). Isolated snRNAs were resolved by denaturing polyacrylamide gel electrophoresis with 7M urea and gel-purified (Supplementary Fig. S1A, S1B). Purified snRNA (2 pmol) was digested with RNase T_1_ (10 U; Thermo Fisher Scientific) in 25 mM ammonium acetate (pH 5.3) or RNase A (10 ng; Thermo Fisher Scientific) in 25 mM ammonium acetate (pH 7.3) at 37 °C for 60 min. The digests were separated at a flow rate of 300 nL/min by capillary LC in a solvent system consisting of 0.4 M 1,1,1,3,3,3-hexafluoro-2-propanol (HFIP) (pH 7.0) (solvent A) and 0.4 M HFIP (pH 7.0) in 50% methanol (solvent B) using a linear gradient (2–100% B in 40 min) (Takakura et al. 2019). The eluent was ionized by an ESI source in negative polarity mode and scanned over an *m/z* range of 600–2000. Data were analyzed on Xcalibur 4.7 software (Thermo Fisher Scientific).

### Pseudouridine quantification by BACS

Pseudouridylation of U2, U4, and U5 snRNAs was detected by BACS (Xu et al. 2024). 2.0 µg total RNA was first treated with 2 U RQ1 DNase (Promega) at 37 °C for 30 min, followed by the addition of 1.0 µL stop solution and incubation at 65 °C for 10 min. RNA was then purified using an RNA Clean & Concentrator (Zymo Research). For 2-bromoacrylamide (2-BAA) derivatization, 1.0 µg DNase-treated RNA was mixed with 20.0 µL 2-BAA buffer (250 mM 2-bromoacrylamide [Enamine], 625 mM potassium phosphate, pH 8.5) and incubated at 85 °C for 30 min, followed by two rounds of purification on Centri-Sep columns (Princeton Separations). For reverse transcription, 400 ng of 2-BAA-treated RNA was mixed with 20 pmol gene-specific primer and 1.0 µL dNTP mix (10 mM each) in Milli-Q water to a total volume of 14.5 µL, heated at 65 °C for 5 min, and chilled on ice. The annealed mixture was combined with 4.0 µL 5× RT buffer, 0.5 µL SUPERase·In RNase inhibitor (20 U/µL; Thermo Fisher Scientific), and 1.0 µL Maxima H reverse transcriptase (200 U/µL; Thermo Fisher Scientific) to a final volume of 20 µL. Reactions were incubated at 50 °C for 30 min and 85 °C for 5 min. PCR was performed in 20 µL reactions containing 2.0 µL cDNA, 4.0 µL 5× Phusion GC buffer, 2.0 µL 2 mM each dNTP, 2.0 µL 30% DMSO, 1.0 µL each of 10 µM forward and reverse primers, and 0.2 µL Phusion DNA polymerase (2 U/µL). Cycling conditions were 98 °C for 30 s; 25 cycles of 98 °C for 10 s, 63 °C for 20 s (59 °C for U4 and U5), and 72 °C for 5 s. To remove residual primers, PCR products were treated with Exonuclease I (New England Biolabs) by adding 2.5 µL 10× Exonuclease I buffer and 2.5 µL Exonuclease I, incubating at 37 °C for 15 min and 80 °C for 15 min, and then purified using an RNA Clean & Concentrator. Nested PCR was carried out with the same reaction composition using 2.0 µL of the first PCR product as template and an annealing temperature of 62 °C, followed by gel purification and Sanger sequencing.

### RNA-sequencing analyses

Mouse liver and testis tissues were harvested immediately following cervical dislocation. Tissues were homogenized in 5 ml TRIzol reagent (Invitrogen #15596026), incubated at 50°C for 5 min, and transferred to a 15-ml tube. Subsequently, 1 ml chloroform was added, vigorously mixed for 30 seconds, incubated on a rotator for 5 min, and centrifuged at 4°C at 12,000 × g for 5 min. The aqueous layer was collected, mixed thoroughly with 2.5 ml isopropanol, and centrifuged again at 4°C at 12,000 × g for 10 min to pellet the RNA. After removing the supernatant, the pellet was washed with 70% ethanol, dried briefly, and dissolved in 1/10 TE buffer. RNA concentrations were quantified by measuring absorbance. After rRNA depletion by mouse/rat riboPOOL RNA-Seq (siTOOLs Biotech, #DP-K024-000055), the remaining RNA was subjected to library preparation with Biorad SEQuoia Express Stranded RNA Library Prep Kit (Bio-Rad, #12017265), according to the manufacturer’s instruction. Deep sequencing was conducted on a NovaSeq X Plus platform (Illumina) with the 150-bp pair-end mode. Raw fastq files were first processed using fastp (ver. 0.24.0) to remove adapter sequences and trim reads to a defined length, using the following parameters:

--umi --umi_loc read2 --umi_len 10 --umi_prefix UMI --detect_adapter_for_pe --trim_front1 2 --trim_front2 0 --max_len1 130 --max_len2 130 --cut_front --cut_tail --cut_window_size 4 -- cut_mean_quality 20 --qualified_quality_phred 15 --length_required 36 --json "${json_report}" --html "${html_report}". Trimmed reads were mapped to the GRCm39 genome using HISAT2 (ver. 2.2.1) with GENCODE vM34 annotation, and sorted BAM files were generated using SAMtools (ver. 1.21). Gene expression levels were quantified from aligned reads using the featureCounts function in Rsubread (ver. 2.20.0) and normalized with DESeq2 (ver. 1.46.0). RNA-seq fastq data have been deposited in the NCBI database under accession number PRJNA1244441.

### Alternative splicing analyses

Alternative splicing was analyzed with two methodologically distinct tools. First, rMATS (v4.3.0; Shen et al. 2014) was applied to the aligned reads as described above. To focus on events with reliable read support, only events with a mean junction-read coverage of at least 10 – computed for each event as the average across replicates of the sum of inclusion- and skipping-junction read counts in each group, averaged across the two groups – were retained for downstream analysis. Second, to provide an orthogonal, isoform-quantification-based assessment, transcripts were quantified with Salmon (v2.0.0) (Patro et al. 2017) using a decoy-aware index built from the GENCODE vM34 mouse transcriptome (GRCm39), with the primary-assembly genome as the decoy sequence. Paired-end reads were quantified with automatic library-type detection and GC-bias correction (salmon quant -l A --gcBias). Local alternative-splicing events were generated from the same GENCODE vM34 annotation using Suppa2 (Trincado et al. 2018), and generateEvents (-f ioe -e SE SS MX RI FL) and PSI values were computed per event with psiPerEvent, and differential PSI between *Coil^7aa^*/*Coil^7aa^* and control mice was tested per tissue with diffSplice -m empirical -gc. SUPPA2 events with non-computable ΔPSI were excluded from subsequent analysis. For comparison between the two tools, gene-level sets of significant and affected calls were derived independently for each tissue using the five event categories common to both tools (SE, RI, MXE/MX, A3SS/A3, A5SS/A5; the AF and AL categories, which rMATS does not output, were excluded). A gene was scored as significant and affected if it harboured at least one event satisfying rMATS FDR < 0.01 or SUPPA2 p < 0.01, together with |ΔPSI| ≥ 0.1. The Venn diagrams in Fig. S4C show the resulting gene sets restricted to the universe of genes testable by both methods (i.e., genes appearing in the post-filter outputs of both tools).

### Preparation of snap-frozen sections

Mice were anesthetized as described above and perfused with 8 ml of 1×HCMF to remove blood. After cervical dislocation, tissues were harvested and rinsed once in 1×HCMF. Gum tragacanth powder (FUJIFILM Wako, #200-02245) was dissolved in water (1 g/ml) and kneaded, then placed onto cork blocks cut to an appropriate size. Tissue samples were placed onto this gum and quickly immersed in isopentane (Nacalai, #26404-75), pre-cooled in liquid nitrogen, for approximately 30 seconds to achieve rapid freezing. The temperature of isopentane was carefully maintained to remain liquid, as excessively cooled isopentane solidifies, impairing proper freezing. Frozen tissue blocks were sectioned at 8 µm using a cryostat at −20°C. Sections were mounted onto poly-L-lysine-coated slides (MATSUNAMI, #S7441) and stored at −80°C until use.

### Immunostaining and histological staining

Frozen sections were fixed in 4% paraformaldehyde (PFA)/HCMF for 20 min at room temperature, washed in 1×PBS, and permeabilized in methanol at −20°C for 5 min. Sections were blocked in 4% skim milk/TBST (TBS (pH 7.5), 50 mM Tris-HCl, 150 mM NaCl, 0.01% Tween20) for 5 min, followed by incubation with primary antibodies diluted in the blocking solution for 1 hour at room temperature. After washing three times in 1×TBST, sections were incubated with fluorescently labeled secondary antibodies for 40 min. Following additional washes with 1×TBST, sections were mounted with polyvinyl alcohol medium containing DABCO 33-LV (Sigma, #290734). Fluorescent images were captured using a CCD camera (DP74; Olympus) attached to a conventional fluorescence microscope (BX51; Olympus) or confocal microscopy (LSM900; Zeiss). 3D rendering images were created using ZEN Blue software (Zeiss). For HE staining, snap-frozen sections were stained with hematoxylin (FUJIFILM Wako, #131-09665) for 5 min, washed under running water for differentiation, then stained with eosin solution (Sigma, #HT110116) for 1 min. Sections were dehydrated sequentially in graded ethanol (75%, 90%, 100%), absolute ethanol, and xylene, and mounted with Entellan New (Sigma #107961), a non-aqueous mounting medium. All the antibodies used in this study are listed in **Supplemental Table 4**.

### Staging of seminiferous tubules

Coarse staging of HE-stained seminiferous tubules (stage I–III, IV–VI, VII/VIII, IX–X, and XI–XII) was performed based on the classical criteria, which consider characteristic cellular and chromatin morphologies (Oakberg 1956). More precise staging and cell type identification were determined as described previously, using a combination of specific staining patterns of PNA (Nakata et al. 2015), gamma Histone H2AX (γH2AX) (Blanco-Rodriguez 2009), DAPI, and coilin. Briefly, each stage was characterized as follows:

Stage I: Pachytene spermatocytes exhibit a large γH2AX focus/domain. Round spermatids are present without distinct dot-like PNA signals. Elongated spermatids display condensed, linear acrosomal signals outlined by PNA staining. Stage II: Pachytene spermatocytes continue to show a prominent γH2AX focus/domain. Round spermatids show Golgi-phase PNA staining adjacent to their nuclei, while elongated spermatids have an expanded PNA signal extending slightly toward the mid-region of the head compared to Stage I. Stage III: Pachytene spermatocytes maintain large γH2AX focus/domain. Round spermatids have broader Golgi-phase PNA signals compared to Stage II. Elongated spermatids exhibit further expanded PNA staining, covering the entire sperm head, with a strong linear signal marking the acrosome. Stage IV: Pachytene spermatocytes display a characteristic γH2AX focus/domain. Round spermatids show cap-phase PNA signals, and elongated spermatids show uniform PNA staining across the entire sperm head. Emergence of cytoplasmic coilin bodies marks the onset of this stage. Stage V: Similar to Stage IV, pachytene spermatocytes and round spermatids show cap-phase PNA signals. However, coilin signals in the cytoplasmic coilin body reach their peak intensity, delineating Stage V. Stage VI: Pachytene spermatocytes and round spermatids with cap-phase PNA signals persist, but coilin signals in the cytoplasmic coilin body start to decrease. Elongated spermatids maintain uniform acrosomal PNA staining. Stage VII: A mixture of preleptotene and pachytene spermatocytes is present; preleptotene cells exhibit diffuse nuclear γH2AX signals, whereas pachytene cells retain large speckles. Round spermatids with cap-phase PNA signals and elongated spermatids aligned linearly along the lumen are observed. Stage VIII: Similar to Stage VII, with elongated spermatids showing more pronounced linear acrosomal PNA signals. Stage IX: Round spermatids are absent, a characteristic shared by stages IX–XII. Acrosome-phase elongated spermatids are present. Spermatocytes are distinguished by their γH2AX patterns. Stage X: Acrosome-phase elongated spermatids are present. Spermatocytes at leptotene and pachytene stages are distinguished by characteristic γH2AX patterns. Stage XI: Acrosome-phase elongated spermatids persist. Spermatocytes predominantly exhibit zygotene and pachytene phases. Diplotene spermatocytes transitioning from late pachytene are also found. Stage XII: Defined by the absence of round spermatids, this stage contains acrosome-phase elongated spermatids and secondary spermatocytes characterized by small, round nuclei. A comprehensive summary of the cellular associations, spermiogenesis steps (Steps 1–16), and specific marker staining patterns across the 12 stages of the seminiferous epithelial cycle is provided in **Supplementary Table 2**.

### Analyses of sperm motility

Mature male mice (>8 weeks old) were euthanized, and epididymides were isolated alongside testes. The cauda epididymis was dissected to remove blood and fat, then briefly immersed in liquid paraffin. After making a small incision in the cauda epididymis, sperm released into droplets were collected and incubated in mHTF medium (Kyudo, Japan) at 37°C in a 5% CO2 incubator for 2 hours. Sperm motility was measured using the Sperm Motility Analysis System (SMAS). Briefly, a 3 µl aliquot of sperm suspension was placed on a sperm analysis slide (sperm plate, DITECT #DP041125, depth 12 µm). Motility parameters (VSL, VCL, VAP, and STR) were quantified automatically by the SMAS system using video recordings (150 fps) from six different microscopic fields per mouse, captured by phase-contrast microscopy equipped with a CCD camera.

## Acknowledgment

We would like to thank Drs Shosei Yoshida and Kiyozumi Daiji for valuable comments and discussions on spermatogenesis, Akira Nakamura for a laboratory space and general discussions, Masahiro Sato for technical advice on micro manipulations, Rei Yoshimoto for advice on rMATS analyses, and Satomi Miyagawa for antibodies used for initial studies. This work is supported by JSPS KAKENHI grant numbers 21H05274, 21H05273, 24H00546, 26H01553, and 26H01553 for S.N., 23K06093 and 16H06279 for H.M., 25K02279 for Y.O., JST-ERATO (JPMJER2002) for T.S., RIKEN TRIP initiative “TRIP-AGIS” for S.I. and Global Facility Center (GFC) at Hokkaido University.

## Figure legends

**Supplemental Fig. S1.**
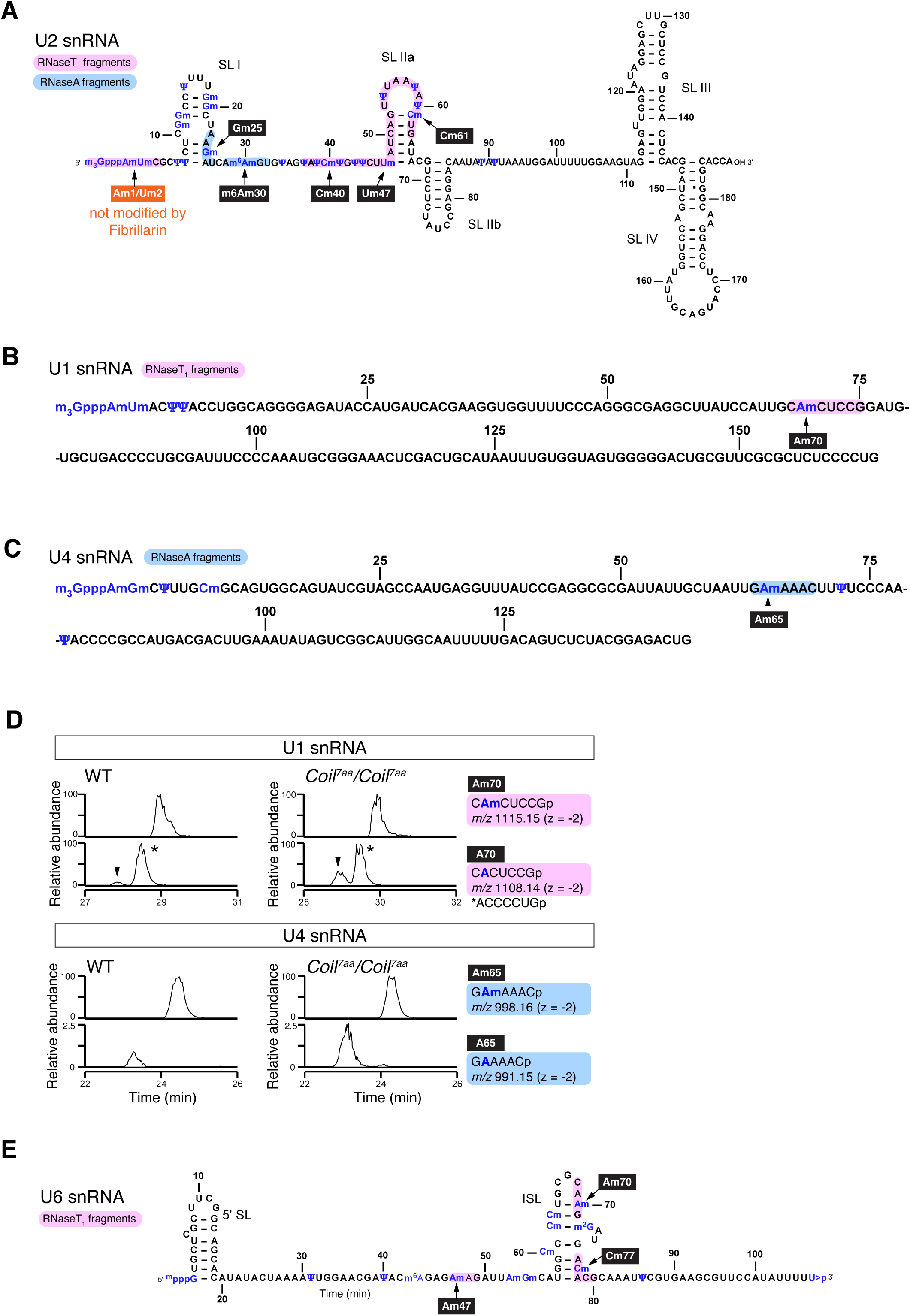
Isolation of snRNAs and LC/MS analysis of U1 and U4 snRNA. (A) Sequence and secondary structure of U2 snRNA. Magenta and light-blue regions indicate RNA fragments analyzed by LC-MS after RNase T_1_ or RNase A digestion, respectively. Positions of modifications were inferred based on previous studies performed on human U2 snRNA (Donmez et al. 2004). Ψ60 was newly identified as a pseudouridine in this study. (B) Sequence of U1 snRNA. The magenta region indicates the RNA fragment analyzed by LC-MS after RNase T_1_ digestion. (C) Sequence of U4 snRNA. The light-blue region indicates the RNA fragment analyzed by LC-MS after RNase A digestion. (D) Extracted-ion chromatograms of RNase T_1_- and A- treated snRNA fragments containing modifications at U1 snRNA Am70 and U4 snRNA Am65. Arrowheads indicate the peaks of fragments containing Am70, and asterisks indicate the peaks derived from another fragment. (E) Sequence and secondary structure of U6 snRNA. Magenta regions indicate RNase T_1_-digested RNA fragments analyzed by LC-MS. Modification sites were annotated based on human studies (Shimba et al. 1995).

**Supplementary Figure S2:**
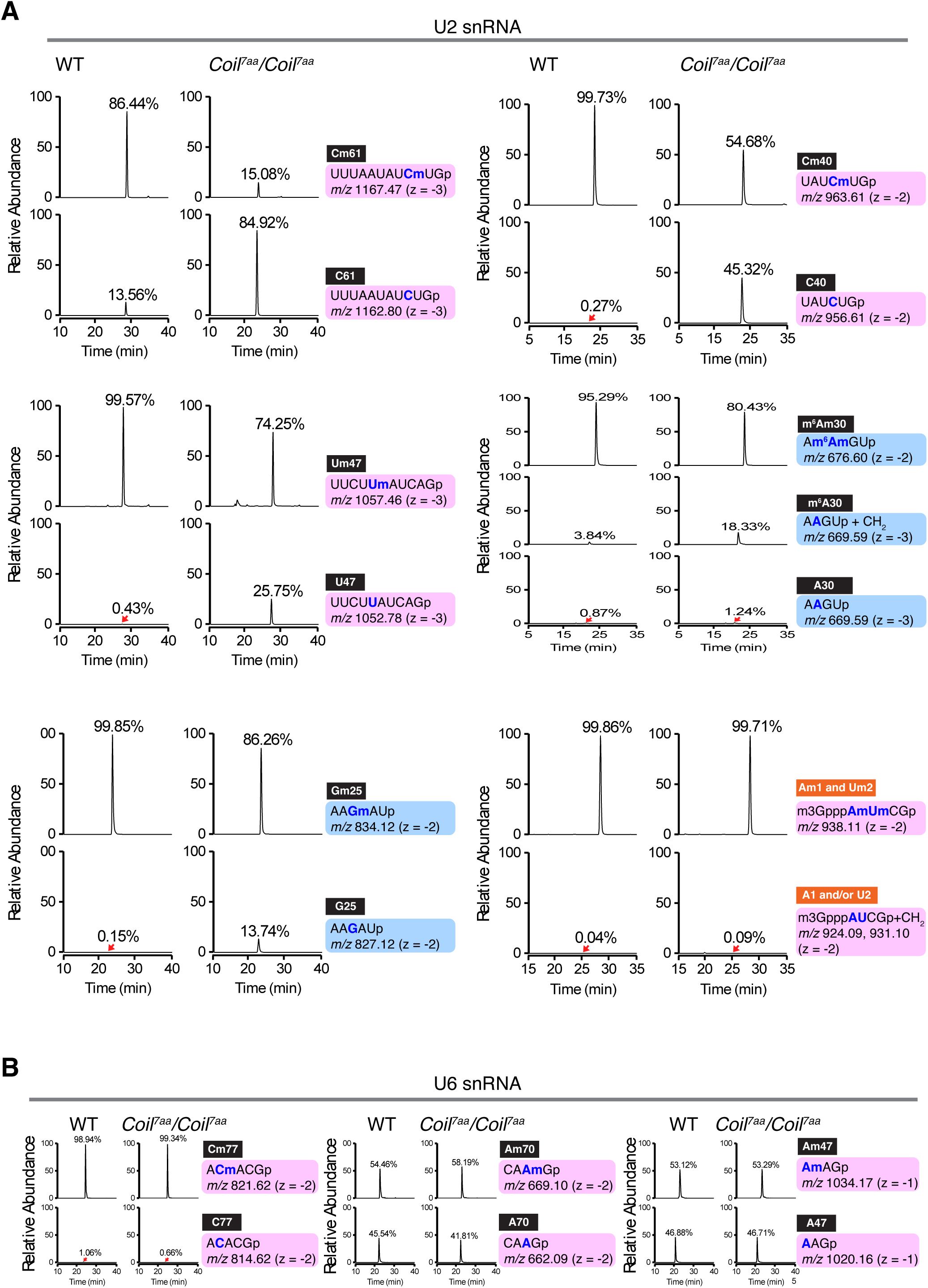
Reproducibility of altered snRNA modifications in an independent biological replicate. (A) Extracted-ion chromatograms of RNase T1- and A-treated U2 snRNA fragments containing modifications Cm61, Cm40, Um47, m6Am30, Gm25, and Am1/Um2, obtained from a separate batch of wild-type and *Coil^7aa^*/*Coil^7aa^* mice. Consistent with the main results in Figure 2, fragments containing unmodified nucleotides increased in *Coil^7aa^*/*Coil^7aa^* mice at all shown positions except the cap Am1/Um2 nucleotides, accompanied by a reduction of fragments with modifications. n.d., not detected. (B) Extracted-ion chromatograms of U6 snRNA fragments containing modifications at Cm57, Am70, and Am47 from the independent replicate. As observed in Figure 2, the ratios of modified versus unmodified fragments remained unchanged between wild-type and *Coil^7aa^*/*Coil^7aa^*mice.

**Supplemental Fig. S3.**
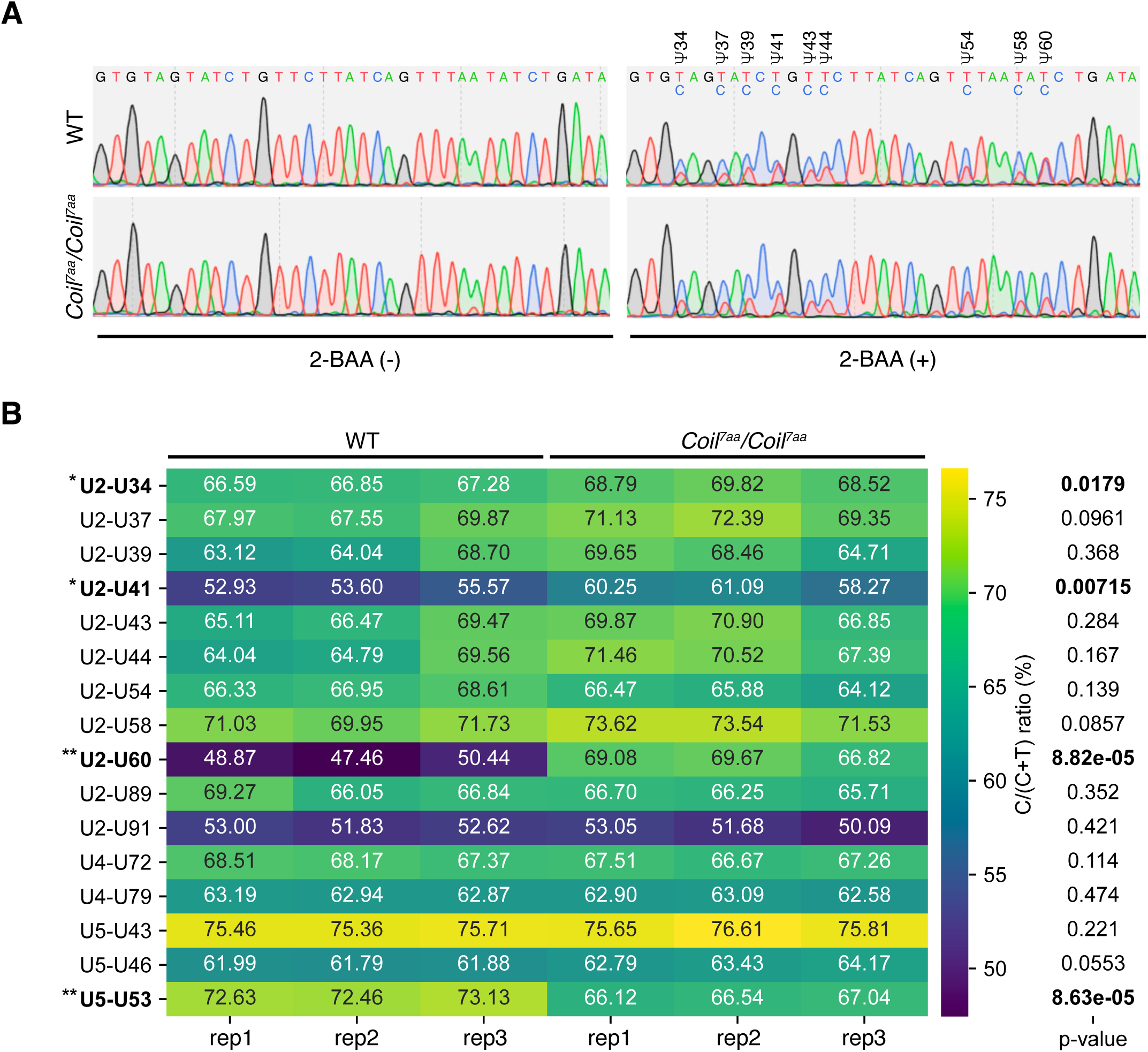
Detection of pseudouridine by BACS. (A) Example Sanger sequencing chromatograms of U2 snRNA between Ψ34 and Ψ60, showing the conversion of the readout from T to C at pseudouridylation sites upon 2-BAA treatment. The positions of the modifications are shown below the chromatogram. (B) Heatmap of T-to-C conversion rates at each detected position on snRNAs. The p-values of the two-sided Student’s T-test between triplicates of WT and KO for each position are shown on the right side. Statistically significant changes were shown in bold.

**Supplemental Fig. S4.**
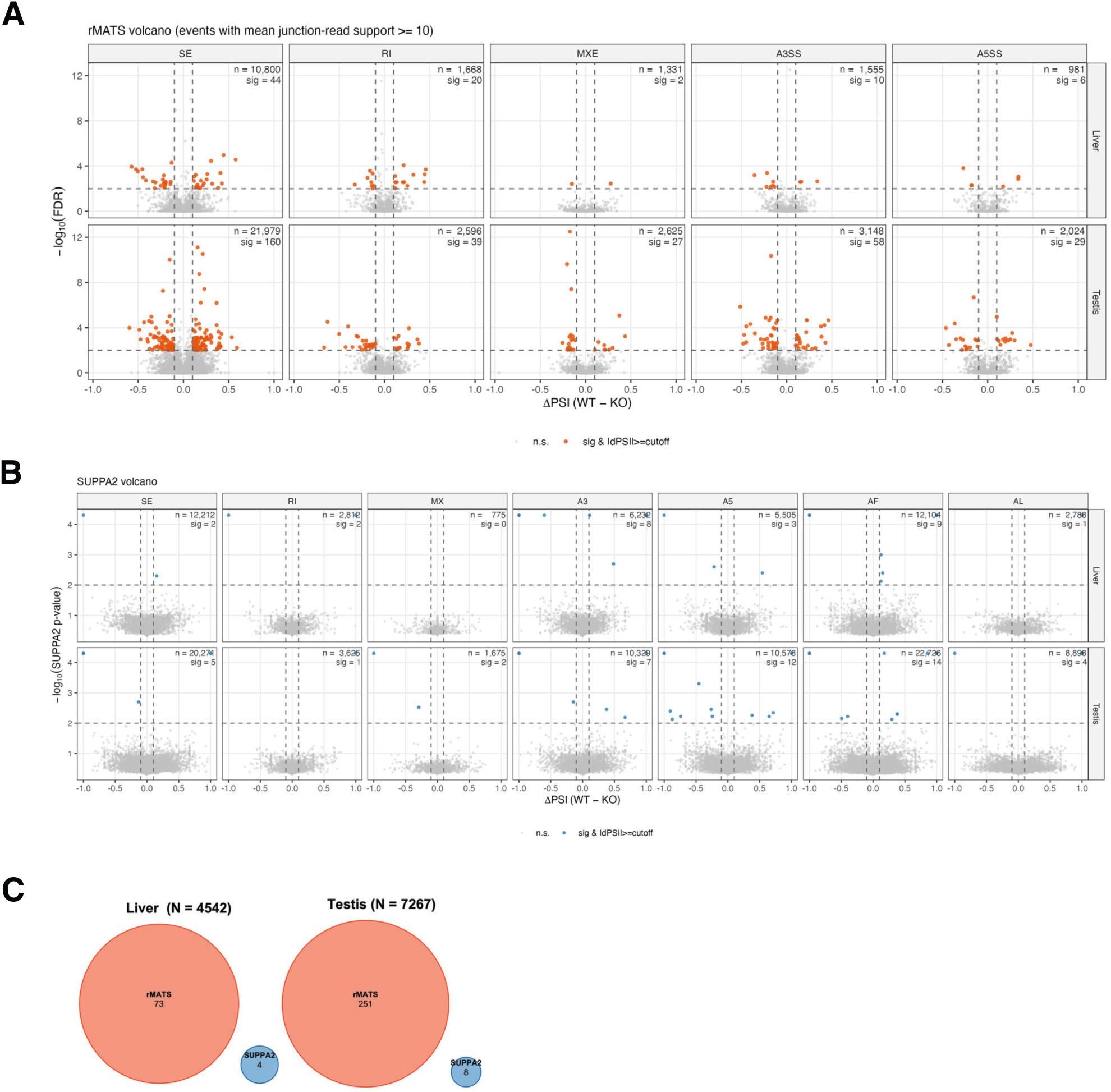
Orthogonal validation of the rMATS alternative-splicing analysis with SUPPA2. (A) Volcano plots of differential splicing called by rMATS in liver (top row) and testis (bottom row), faceted by event type (SE, skipped exon; RI, retained intron; MXE, mutually exclusive exons; A3SS, alternative 3′ splice site; A5SS, alternative 5′ splice site). Only events with a mean junction-read coverage of ≥10 across all replicates are shown. Grey, n.s.; blue, significant (FDR < 0.01) but subtle (|ΔPSI| < 0.1); orange, significant (FDR < 0.01) with |ΔPSI| ≥ 0.1. Dashed lines mark FDR = 0.01 (horizontal) and |ΔPSI| = 0.1 (vertical). Numbers in each facet give the total number of events plotted (n) and the number of significant events (sig). (B) Volcano plots of differential splicing called by SUPPA2 on the same liver and testis datasets, faceted by event type and tissue as in (A). Events whose ΔPSI was not computable (nan, reflecting insufficient isoform-level TPM support in at least one condition) or whose p-value was exactly 1.0 are excluded. Thresholds and color scheme as in (A). (C) Area-proportional Venn diagrams of the gene-level "significant and affected" sets identified by rMATS and SUPPA2 in each tissue. A gene was scored as significant and affected if it harboured at least one event meeting the joint criterion of statistical significance (rMATS FDR < 0.01 or SUPPA2 p < 0.01) and effect size (|ΔPSI| ≥ 0.1), considering the five event categories common to both tools (SE, RI, MX/MXE, A3/A3SS, A5/A5SS). Gene sets are restricted to the universe of genes testable by both methods after the read-support filters described in (A) and (B); N indicated in each panel title. No gene was identified concordantly by the two methods in either tissue.

**Supplemental Table 1.** Percentage of modified bases in the liver of WT and Coil KO mice

**Supplemental Table 2.** Counts of mapped reads used for RNA-Seq analyses

**Supplemental Table 3.** Summary of cellular associations, spermiogenesis steps, and specific markers across the seminiferous epithelial cycle

**Supplemental Table 4.** Primers and antibodies used in this study

